# Deep *in vivo* brain metabolomics maps vulnerability-associated and cocaine-induced neurochemical states

**DOI:** 10.64898/2026.08.14.744954

**Authors:** Caitlin N. Cain, Pavlo Popov, Nicholas M. Oliver, Juan C. Vázquez López, Charles R. Evans, Elaine K. Hebda-Bauer, Huda Akil, Robert T. Kennedy

## Abstract

Innate temperament affects vulnerability to substance use disorders (SUDs). Identifying the neurochemistry that differentiates temperaments, alongside the metabolic impact of drugs of abuse, can provide new insights into pathways to SUD and treatments. We show that heritable emotional temperaments associated with SUD vulnerability are encoded by distinct extracellular neurochemical states in the living brain. We apply untargeted metabolomics to microdialysis samples to define the chemical composition of the living brain metabolome with unprecedented depth. Using two phenotypes from a rodent model of diverging temperaments linked to SUDs, dialysate samples were collected before and after acute exposure to cocaine (15 mg/kg, i.p.). Metabolomic analysis revealed previously unknown basal differences in stress-response, bioenergetics, and amino acid neurotransmitters and their related pathways between the two phenotypes. Cocaine administration also had a profound impact on the metabolome, altering amino acid, purine, and arachidonic acid pathways along with cognition-related metabolites. Despite these broad metabolomic changes, cocaine administration narrowed several pre-existing differences between phenotypes, highlighting divergent responses across genetic lines. These findings demonstrate that *in vivo* metabolomics not only offers novel insights into the neurochemistry underlying temperament and drug response but also serves as framework to advance a systems-level understanding of other brain-related disorders.

## INTRODUCTION

Substance use disorders (SUDs) are chronic, relapsing conditions characterized by the compulsive use of addictive substances despite their long-term adverse health outcomes.^1^ Most drugs of abuse activate the brain’s reward circuitry, reinforcing drug-taking behavior and impairing overall function.^2,3^ However, vulnerability to SUDs extends beyond these neurobiological effects to also include genetic^4,5^ and environmental^6,7^ influences. For instance, susceptibility to substance use is closely linked to heritable traits such as emotional temperament. Human genetic studies have demonstrated that both externalizing traits (e.g., heightened novelty-seeking, risk-taking, impulsiveness) and internalizing traits (e.g., anxiety, depression) can contribute to SUD risk.^8–11^ However, a critical gap remains in understanding how these genetic traits translate into the neurochemistry and behaviors that drive addiction. We leverage untargeted metabolomics to capture the *in vivo* neurochemical dynamics that define these heritable emotional temperaments under baseline and psychostimulant-induced conditions. We demonstrate that novel stress-related, bioenergetic, and amino acid metabolites characterize these temperament differences. Acute cocaine exposure induces large shifts across the brain metabolome, disrupting several pathways crucial for cognitive function and reducing some phenotypic differences.

The brain metabolome includes signaling molecules, such as small molecule and lipid neurotransmitters and neuromodulators, as well as compounds involved in energy metabolism, neuroprotection, plasticity, and cell growth. Untargeted metabolomics of tissue has been used to generate snapshots of brain chemical composition, capturing the interplay between genetic and environmental interactions and provide insight into biochemistry underlying brain function.^12–21^ Ding et al. recently established a foundational resource for brain metabolomics by annotating 1,547 unique metabolites across 10 brain regions in aging mice.^15^ In the context of SUDs, *ex vivo* metabolomic studies have revealed alterations in neurotransmission and energy metabolism with drug exposure.^22–26^ However, endpoint tissue analyses cannot capture the dynamic metabolic changes linked to behavior, and the isolation process risks altering metabolite levels.^27^ In addition, it is not feasible to differentiate intracellular and extracellular metabolites, obscuring signaling dynamics. In contrast, *in vivo* sampling methods, such microdialysis and solid-phase microextraction, preserve the native biochemical environment and enable capturing extracellular signaling events that directly reflect ongoing neural activity, behavior, and pharmacological response. Coupling such sampling with metabolomics offers an opportunity to link neurochemistry with functional outcomes in ways not accessible through *ex vivo* analyses.

Initial *in vivo* untargeted brain metabolomic studies relying on liquid chromatography-tandem mass spectrometry (LC-MS/MS) for analysis have been able to identify up to 120 compounds. These studies have examined neurochemical changes associated with fluoxetine administration,^28^ deep brain stimulation,^29^ traumatic brain injury,^30,31^ and sleep states.^32,33^ Despite promising results, a vast portion of the brain extracellular metabolome remains uncharacterized primarily due to low physiological concentrations, sample volumes, and/or instrument sensitivity. We recently expanded compound identification efforts for dialysate by optimizing LC-MS/MS acquisition parameters, sample concentration, and run times.^34,35^ In total, we identified approximately 500 compounds in dialysate from the rat striatum,^34,35^ demonstrating the chemical complexity of brain extracellular space.

Leveraging the chemical richness provided by coupling *in vivo* microdialysis with LC-MS/MS, this study investigates the brain extracellular metabolome associated with different temperaments and in response to cocaine administration. We use an animal model of heritable emotional temperament, selectively-bred high responder (bHR) and low responder (bLR) rats, to elucidate how genetic vulnerability to SUDs alters the brain metabolome. In outbred rats, high exploratory locomotion (i.e., HRs) predicts a propensity for psychostimulant self-administration and other externalizing behaviors, whereas lower exploratory responses (i.e., LRs) are associated with anxiety-like and internalizing behaviors.^36,37^ Selective breeding for novelty-induced locomotion has produced a well-validated animal model for studying the heritable temperaments implicated in SUDs.^38,39^ Gene expression studies have shown that these animals are differentiated by their dopaminergic,^40,41^ stress,^37,42^ fibroblast growth factor,^41,43–47^ and energy^48^ systems at both baseline and in response to drugs of abuse. Yet little is known about the metabolomic differences among the bHR/bLR phenotypes. Focusing on the nucleus accumbens, a brain region central to processing rewards and mediating the addictive effects of drugs, we previously found significant phenotype-based differences in basal and cocaine-evoked extracellular dopamine and norepinephrine.^49^ Building upon this work, we use untargeted metabolomics to discover the neurochemical profiles underlying different temperaments and the subsequent alterations induced by acute drug exposure *in vivo*. Our findings identify links between neurochemistry and temperament as well as acute metabolome rearrangements evoked by cocaine. Integrating *in vivo* metabolomics into broader multi-omic frameworks will help advance a systems-level understanding of brain-related disorders.

## RESULTS

### Metabolomic study overview and data quality assessment

Fig. 1A depicts our study design to explore the brain extracellular metabolome at baseline and in response to cocaine. Microdialysis probes were placed to sample from the rostral dorsomedial striatum, the nucleus accumbens core, and the lateral nucleus accumbens shell (Fig. S1). A longer probe membrane was selected to maximize relative metabolite recovery while sampling across these interconnected reward-processing brain regions. Our experiment included male and female bHR and bLR rats (*N* = 8 rats/group), whose selectively bred phenotypes were confirmed by testing for exploratory locomotor response to a novel environment. A two-way analysis of variance (ANOVA) revealed a significant main effect of bHR/bLR phenotype on locomotor response (*F*_1,28_ = 1636.5, *p* < 0.0001), with no significant effects of sex or a sex-phenotype interaction (Fig. 1B). After 5 days of surgical recovery, dialysate samples from these regions were collected during a three-hour baseline period and for three hours after cocaine administration (15 mg/kg, i.p.). All dialysate samples were then analyzed using full-scan LC-MS. To provide deep metabolome coverage, our assay used both reversed-phase (RPLC) and hydrophilic interaction LC (HILIC) as well as positive and negative MS ionization modes. As illustrated by the chromatograms in Fig. S2, this workflow generated four data sets (one for each LC-MS combination) to identify baseline and cocaine-induced metabolomic differences.

**Figure 1.**
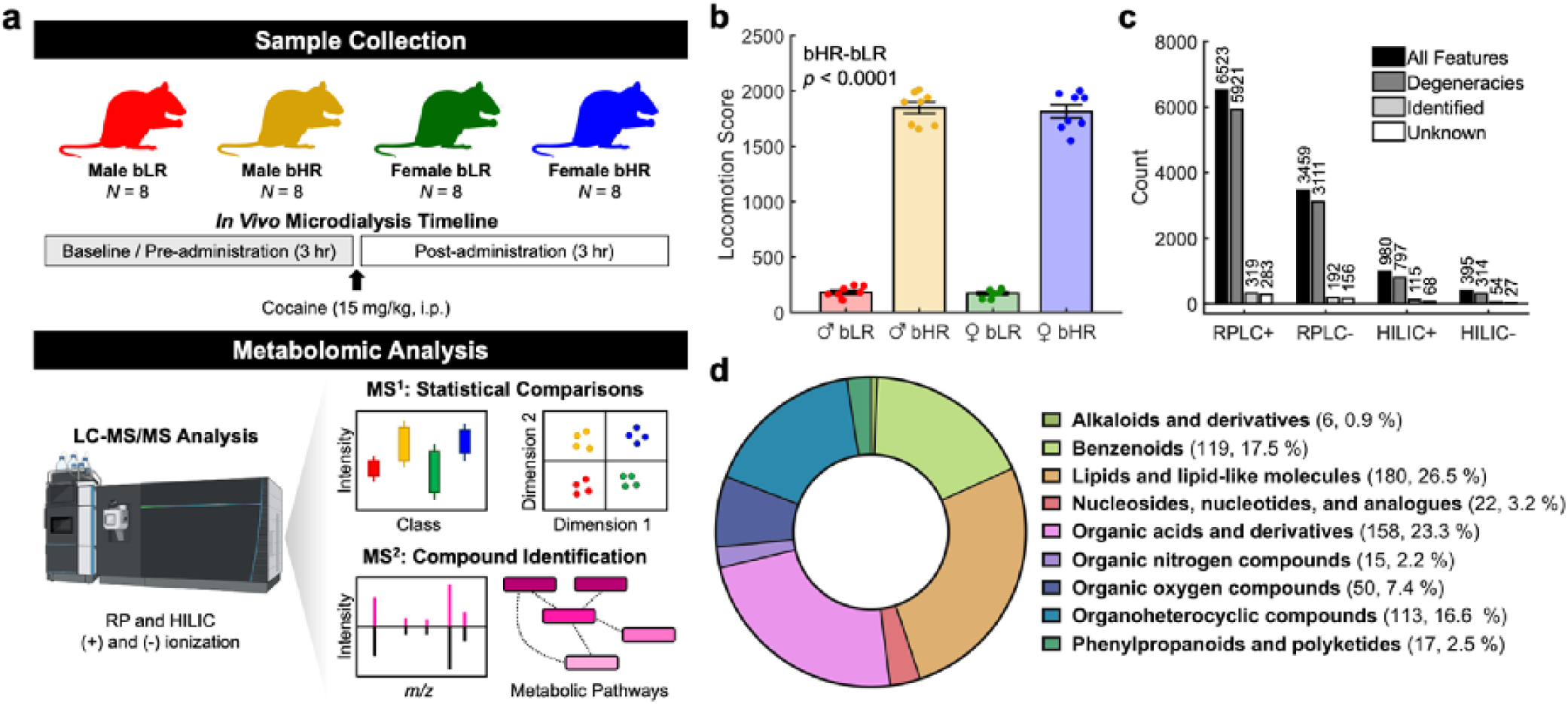
Overview of the experimental design and data set. **a)** Graphical illustration of the experimental workflow to acquire the *in vivo* brain metabolome data. **b)** Difference in locomotor response to a novel environment between bHR and bLR rats. **c)** Number of features, degeneracies, compound identifications, and unknown “feature groups” made in each chromatographic data set. **d)** Chemical composition of the metabolites identified in the brain metabolomic samples. Classifications were made using the ClassyFire categories.

Ion features (i.e., unique chromatographic signals at a specific retention time and *m/z*) among all dialysate samples for each LC-MS data set were first detected and aligned using tile-based Fisher ratio (F-ratio) analysis.^50^ The number of ion features detected ranged from 395 for the HILIC-data set to 5,223 for RPLC+ data set (Fig. 1C). Ion features arising from the same analyte (e.g., adducts, isotopes, and in-source fragments) were then grouped together using lack-of-fit (*LOF*) clustering.^35^ Depending on the chromatographic data set, these degenerate ion features constituted ∼80-91 % of the total number of ion features detected (Fig. 1C). To identify the compounds associated with *LOF* feature groups, tandem mass spectra (MS/MS) were acquired using iterative precursor exclusion on a 10× preconcentrated pooled dialysate sample. A total of 680 unique metabolites were annotated using public databases and our previously published neurochemical library,^34^ according to the spectral search criteria outlined in the Methods section and in accordance with the Metabolomics Standards Initiative.^51^ The remaining 534 “feature groups” were unidentified. The identified metabolites were then categorized into nine chemical superclasses by the ClassyFire system (Fig. 1D). As highlighted in previous work,^34^ the most common compound classes in the brain extracellular metabolome are lipid and lipid-like molecules (26.6%), organic acids and derivatives (23.3%), benzenoids (17.6%), and organoheterocyclic compounds (16.7%). The remaining ∼16% of the metabolome was classified as alkaloids, nucleosides, nucleotides, organic nitrogen and oxygen compounds, phenylpropanoids, and polyketides.

Signal performance and reproducibility were monitored during LC-MS/MS analyses using a pooled dialysate quality control (QC) reference sample that was repeatedly injected throughout the sequence. Principal component analysis (PCA) of the annotated metabolites demonstrated that the QC injections clustered tightly together, indicating minimal technical variation (Fig. S3A). Furthermore, 90.1% of the metabolites identified in the QC reference injections exhibited a relative standard deviation (RSD) of less than 20% (Fig. S3B). These findings highlight the reproducibility and technical quality of the metabolomic data set.

### Baseline metabolomic differences across sex and phenotype

Baseline dialysate samples were first compared to determine if there were significant differences between the neurochemical profiles between sexes or behavioral phenotype (bHR vs. bLR). A partial least squares-discriminant analysis (PLS-DA) model showed clustering according to both sex and phenotype (Fig. 2A). Permutational multivariate analysis of variance (PERMANOVA) was subsequently performed on the data to statistically verify the differences between sex and phenotypes. PERMANOVA confirmed significant effects of both sex (*R*^2^ = 0.07; *p* = 0.03) and phenotype (*R*^2^ = 0.07; *p* = 0.03) on the baseline metabolome. Two-way ANOVAs were performed for each metabolite to assess the effects of phenotype, sex, and their interaction. Metabolites were deemed statistically significant for main effects of sex or phenotype, following Benjamini-Hochberg false discovery rate (FDR) correction (*q* < 0.1). In total, 81 metabolites were uniquely significant for sex, 39 metabolites were uniquely significant for phenotype, and 25 metabolites were shared between both main effects (Fig. 2B). Lists of these compounds are provided in Table S1-S2. Herein, we will focus our analysis on the metabolites that were significantly different between the bHR and bLR phenotypes (see Fig. S4 for sex differences). Based on the volcano plot in Fig. 2C, most metabolites exhibiting phenotype-based differences had higher signals in the dialysate samples from bHRs. Pathway enrichment analysis was performed using the metabolites with a statistical difference in bHRs and bLRs. While some metabolites identified have yet to be mapped to biological origins,^34,52^ Fig. 2D demonstrates that several metabolic pathways are implicated in bHR/bLR differences. These altered pathways include alanine, aspartate, and glutamate metabolism; glycine, serine, and threonine metabolism; valine, leucine, and isoleucine biosynthesis; arginine biosynthesis; tryptophan metabolism; and tricarboxylic acid (TCA) cycle.

**Figure 2.**
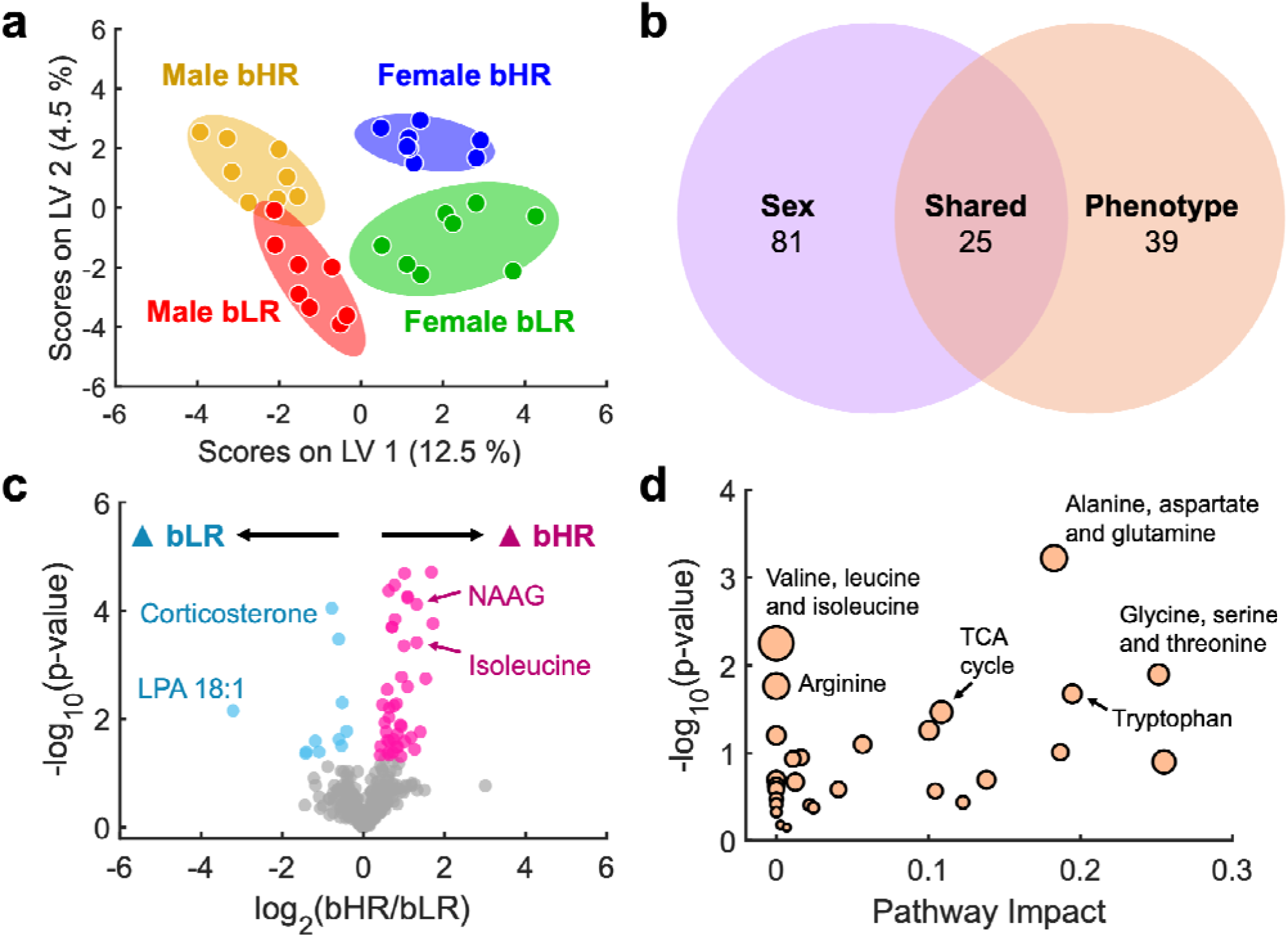
Overview of the baseline neurochemical signature of the bLR and bHR rodent model. **a)** Partial least squares-discriminant analysis (PLS-DA) of the baseline metabolomic samples. Ellipses represent the 95 % confidence interval. Each phenotype-sex combination (male bLRs = red; male b Rs = yellow; female bLRs = green; female bHRs = blue) is tightly clustered together. **b)** Venn diagram depicting the number of unique metabolites with a significant effect for sex (purple), significant effect for phenotype (orange), and the metabolites exhibiting significance for both main effects (overlapped area; “shared”). **c)** Volcano plot of metabolites differentiating bLR and bHR rats. Gray points indicate non-significant features; teal and pink points denote metabolites enriched in bLRs and bHRs, respectively (selected via two-way ANOVA with FDR correction, *q* < 0.1; raw *p*-values displayed). **d)** Pathway analysis of the metabolites that differed between the phenotypes.

Building on prior transcriptomic^37,42^ and behavioral (e.g., Fig. 1B) evidence of altered stress systems between bHRs and bLRs, we first screened the list of phenotype-defining metabolites for potential stress-related compounds. Three candidates were identified: corticosterone, 1-oleoyl lysophosphatidic acid (LPA 18:1), and arachidonoyl ethanolamide (Fig. 3A-C). Corticosterone serves as the primary stress hormone in rodents, whereas LPA 18:1^53–55^ and arachidonoyl ethanolamide^56,57^ are associated with emotional regulation. Each metabolite displayed a variable importance in projection score greater than one (VIP > 1) on the second latent variable (LV 2) of the PLS-DA model, underscoring their contribution to the observed separation of dialysate samples based on phenotype. To confirm the identities for these molecules, their retention times were matched to analytical standards (left panels in Fig. 3A-C).^34^ MS/MS spectra from the dialysate samples were highly consistent with analytical standards, with dot product (DP) and entropy scores (ES) ≥ 700 and 0.65, respectively (middle panels in Fig. 3A-C).^34,35^ Using the precursor ion that triggered the MS/MS event, intensities from each baseline dialysate sample were quantified (right panels in Fig. 3A-C). A two-way ANOVA showed that these three compounds only had a significant effect for phenotype. Both corticosterone (*F*_1,28_ = 19.46, *p* = 0.0001) and LPA 18:1 (*F*_1,28_ = 19.52, *p* = 0.0001) were elevated in bLRs. Meanwhile, arachidonoyl ethanolamide was significantly higher in bHRs (*F*_1,28_ = 15.07, *p* = 0.0006).

**Figure 3.**
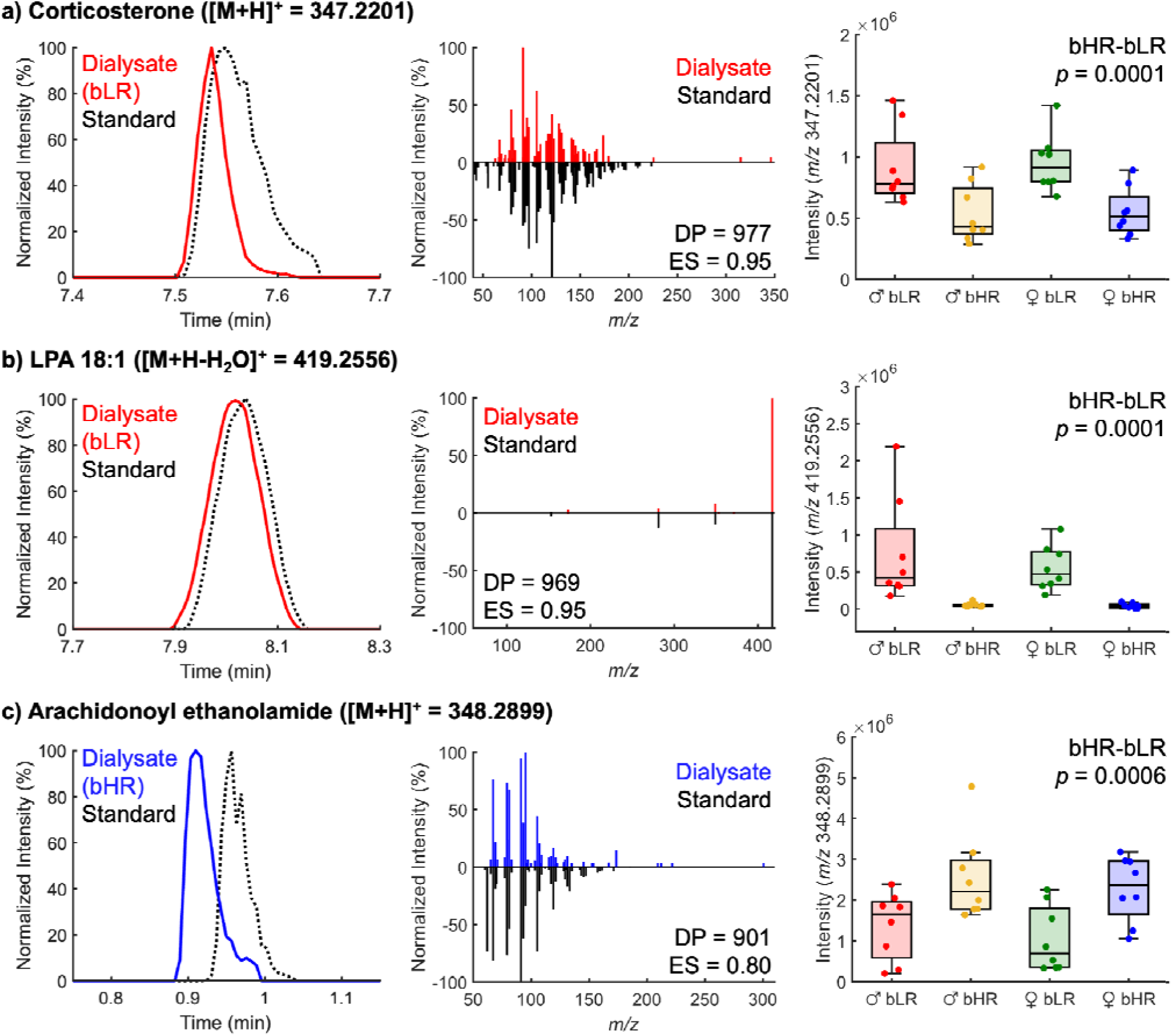
Identification of stress-related compounds in brain dialysate. Chromatographic and mass spectrometry data for **a)** corticosterone, **b)** 1-oleoyl lysophosphatidic acid (LPA 18:1), and **c)** arachidonoyl ethanolamide. Within each panel: Left) Extracted ion chromatograms showing the alignment of the metabolite in the dialysate (solid colored line) with its corresponding standard (dashed black line). Middle) MS/MS reflection plots showing the head-to-tail comparison of the experimental fragmentation pattern (colored) against the reference standard spectrum (black). Dot product (DP) and entropy scores (ES) are provided. Right) Box plots of the relative intensity measured for each compound in each phenotype-sex combination. The central line on the box plot represents the median while the whiskers on the box plots represent the interquartile range (IQR). Each dot represents data from a single animal (*N* = 8 rats/group). Calculated *p*-values for the main effect of phenotype are also indicated.

Motivated by prior transcriptomic work that fatty acid oxidation and TCA cycle function was upregulated in bLRs,^48^ we also investigated several bioenergetic metabolites that were found to differ with phenotype. For example, acylcarnitines act as essential shuttles that transport fatty acids across the mitochondrial membrane for energy production via β-oxidation.^58^ We identified six short- to medium-chain acylcarnitines and two dicarboxylic acylcarnitines in our dialysate samples, all of which exhibited significant main effects for both sex and phenotype (Table S3). Across both sexes, bHRs consistently maintained higher extracellular levels than bLRs (Fig. 4A). Two-way ANOVA also revealed significant sex × phenotype interactions for propionylcarnitine (C3), butyrylcarnitine (C4), and adipoylcarnitine (C6-DC), reflecting a phenotypic divergence that was more pronounced in males than females (Fig. 4A). Aligning with previous transcriptomic findings,^48^ these extracellular acylcarnitine patterns correspond to potential alterations in fatty acid utilization between phenotypes. Further supporting these differences in bioenergetics, we also found two products of incomplete lipid catabolism,^59^ 3-hexendioic acid (*F*_1,28_ = 33.48, *p* < 0.0001) and 3-hydroxyoctanoic acid (*F*_1,28_ = 25.66, *p* < 0.0001), were significantly higher in bHR animals (Fig. 4B). Lastly, both transcriptomic evidence^48^ and the pathway analysis in Fig. 2D indicated differences in TCA cycle function between the two phenotypes. Fig. 4C shows that extracellular levels for two intermediates in the TCA cycle, aconitate (*F*_1,28_ = 19.05, *p* = 0.0002) and α-ketoglutarate (*F*_1,28_ = 39.14, *p* < 0.0001), were significantly lower in bLRs.

**Figure 4.**
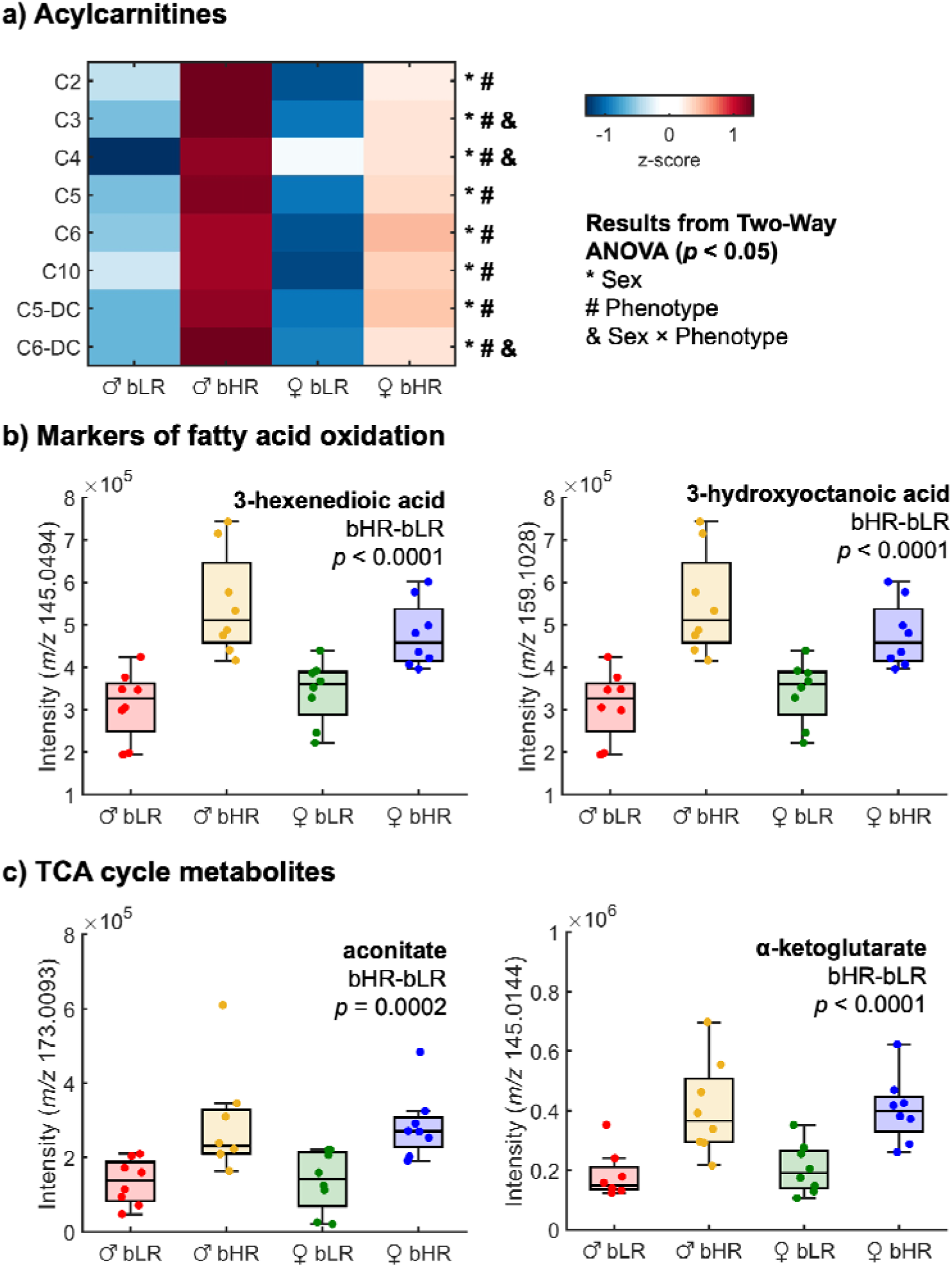
Baseline phenotypic differences in energy metabolism. **a)** Heatmap illustrating the average intensity of identified acylcarnitines across experimental groups. Statistical significance was determined via two-way ANOVA, with symbols denoting a significant main effect of sex (*), a significant main effect of phenotype (#), and a significant sex × phenotype interaction (&). The color scale ranges from −1.5 to +1.5, where color represents the z-score of the metabolite intensities. **b-c)** Box plots of the relative intensity measured for two markers of mitochondrial fatty acid oxidation and two TCA metabolites. Box plots are shown with center line as median, extending to the IQR, with whiskers to the highest and lowest non-outlier points. Each dot represents data from a single animal (*N* = 8 rats/group). Calculated *p*-values for the main effect of phenotype are also indicated.

The pathway analysis in Fig. 2D also highlighted several differences in amino acid metabolism between the bHR and bLR phenotypes. Relating to alanine, aspartate, and glutamate metabolism, significant phenotype-based differences were found for *N*-acetylaspartate (*F*_1,28_ = 18.84, *p* = 0.0002), *N*-acetylaspartylglutamate (NAAG; *F*_1,28_ = 21.17, *p* < 0.0001), and asparagine (*F*_1,28_ = 25.64, *p* < 0.0001; Fig. 5A). Within arginine biosynthesis, we found significant differences in arginine (*F*_1,28_ = 56.87, *p* < 0.0001) and *N*-acetylornithine (*F*_1,28_ = 40.18, *p* < 0.0001; Fig. 5B). Both pathways in Fig. 5A-B also interface with the TCA cycle described earlier in Fig. 4B, where α-ketoglutarate serves as the key metabolite linking neurotransmission and energy production. Similarly, two analytes related to glycine, serine, and threonine metabolism had a main effect of phenotype (Fig. 5C): serine (*F*_1,28_ = 22.77, *p* < 0.0001) and creatine (*F*_1,28_ = 21.38, *p* < 0.0001). As illustrated in Fig. 5D, significant phenotype-based differences were observed for isoleucine (*F*_1,28_ = 15.61, *p* = 0.0005) and its related branched-chain amino acid metabolite, 3-methyl-2-oxopentanoic acid (*F*_1,28_ = 20.29, *p* = 0.0001). Lastly, Fig. 5E shows three compounds related to tryptophan metabolism that have significant phenotype differences: tryptophan (*F*_1,28_ = 13.87, *p* = 0.0009), 2-oxoadipate (*F*_1,28_ = 15.12, *p* = 0.0006), and 3-hydroxyanthranilate (*F*_1,28_ = 25.38, *p* < 0.0001). The overall elevation of these amino acid metabolites in bHRs compared to their bLR counterparts reflect differences in their energy production and neurotransmission.

**Figure 5.**
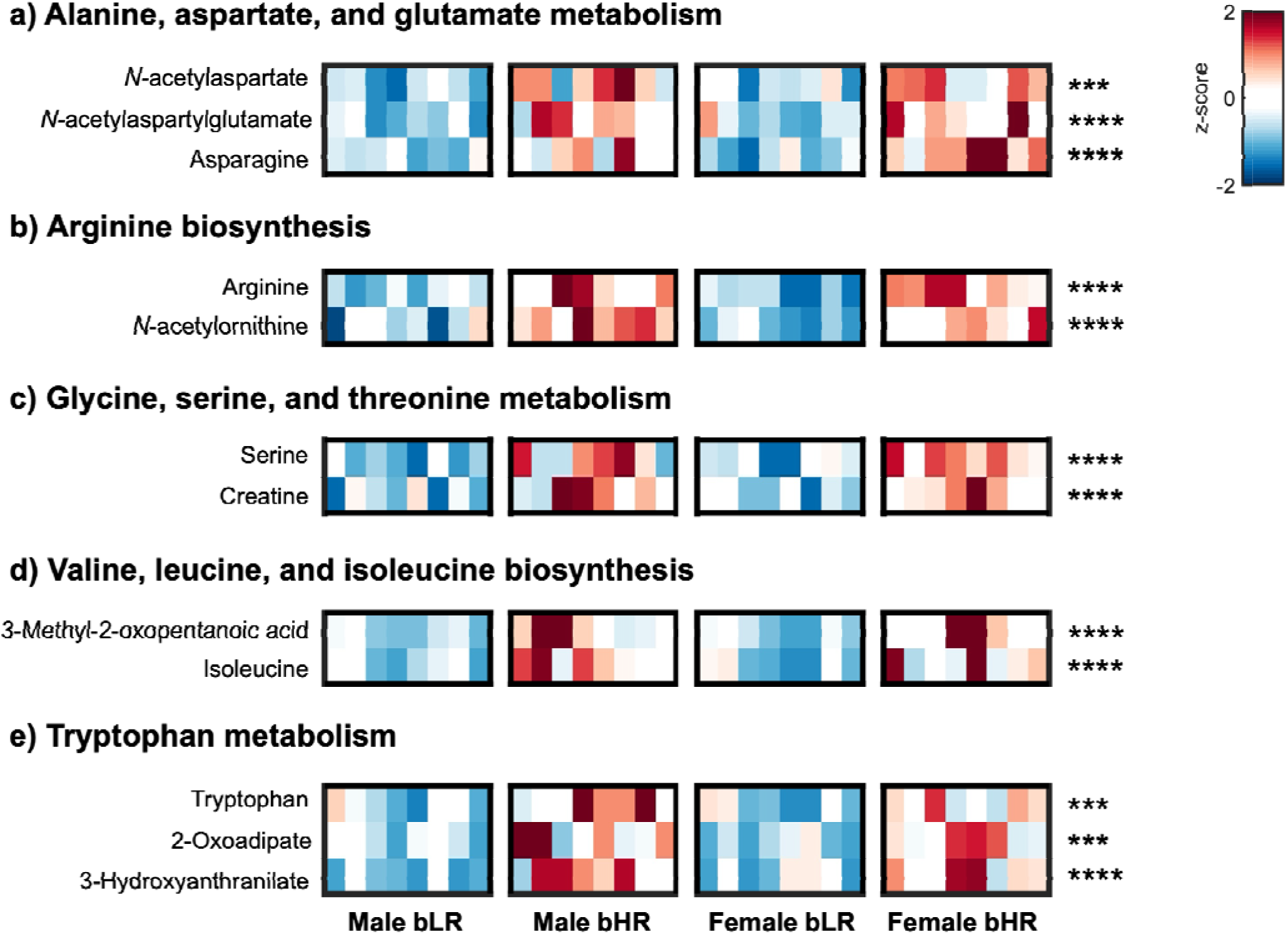
Baseline phenotype differences in amino acid metabolism. Heatmaps of metabolite intensities across phenotype-sex groups. Each square within a phenotype-sex cohort represents an individual animal. Metabolites are categorized by their assigned pathway: **a)** alanine, aspartate, and glutamate metabolism; **b)** arginine biosynthesis; **c)** glycine, serine, and threonine metabolism; **d)** valine, leucine, isoleucine biosynthesis; **e)** tryptophan metabolism. The color scale ranges from −2.0 to +2.0, where color represents the z-score of the metabolite intensities. Asterisks indicate the main effect of phenotype from a two-way ANOVA: *** *p* < 0.001, **** *p* < 0.0001.

### Effect of cocaine on the brain metabolome

Next, we characterized the brain metabolome transition from baseline to a cocaine-induced state (15 mg/kg, i.p.) across the bHR and bLR phenotypes. The PLS-DA scores plot in Fig. 6A visualizes the differences in the neurochemical profiles collected pre- and post-cocaine administration. The separation of these dialysate samples along LV 1 is strongly driven by the metabolomic response to cocaine administration, whereas LV 2 and LV 3 primarily reflect the sex- and phenotype-based differences, respectively. PERMANOVA results confirmed that although cocaine administration was the primary driver of metabolomic differences in these dialysate samples (*R*^2^ = 0.1; *p* = 0.001), sex (*R*^2^ = 0.06; *p* = 0.01) and phenotype (*R*^2^ = 0.04; *p* = 0.03) remained significant contributors. To test whether baseline phenotypic distinctions persist following acute drug exposure, an additional PLS-DA model was built solely with the post-cocaine dialysate samples (Fig. S5A). Interestingly, while post-cocaine female bHR and bLR metabolomes converged toward a shared state, male bHR and bLR metabolomes remained separated (Fig. S5A). Sex-specific volcano plots confirmed this observation, revealing that acute drug exposure eliminated most baseline phenotypic differences in females but not in males (Fig. S5B-C). These results highlight a complex interaction of sex, phenotype, and drug effects that shape the brain’s metabolomic response to acute cocaine exposure.

**Figure 6.**
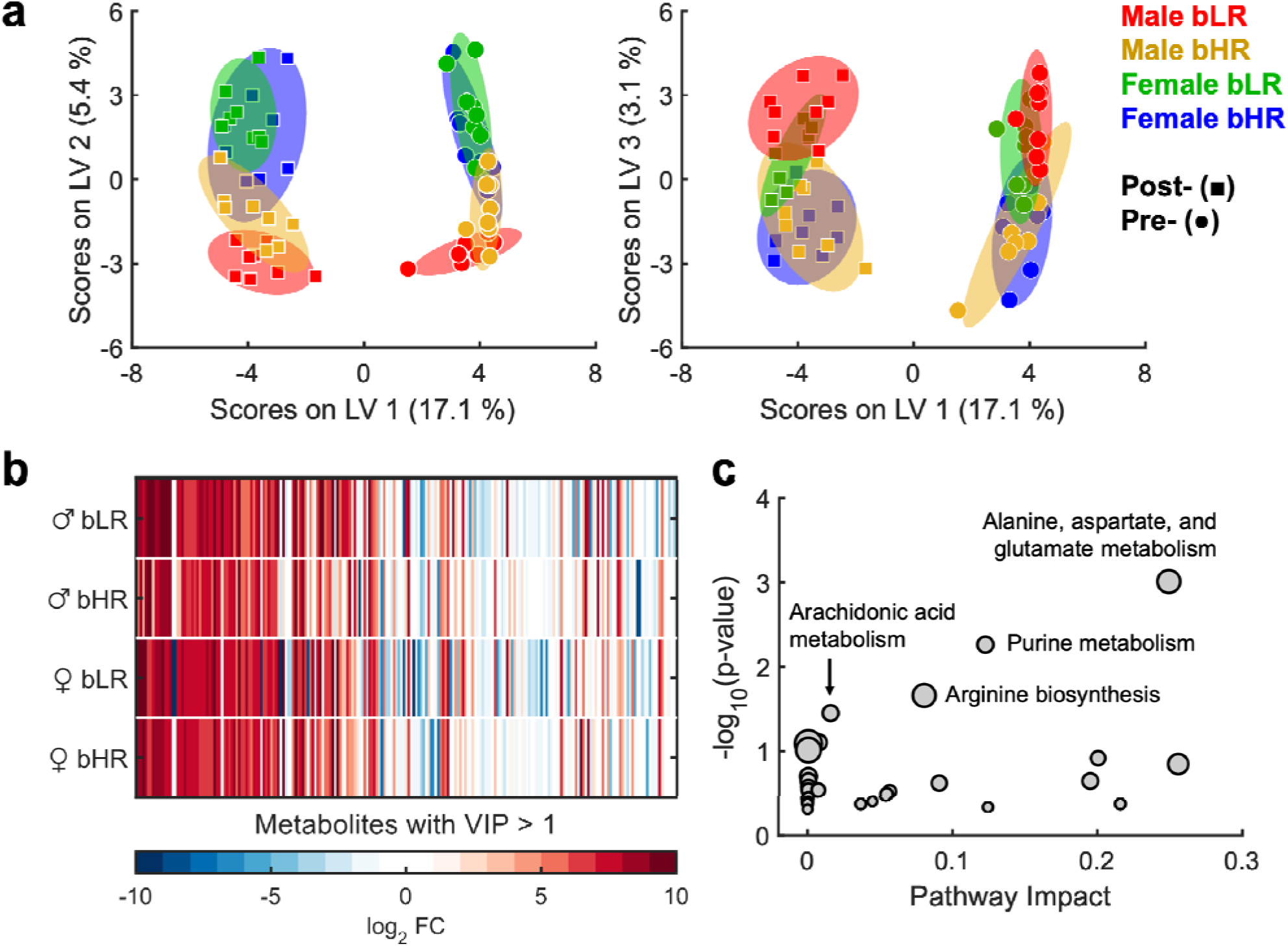
Acute cocaine exposure impacts the living brain metabolome. **a)** PLS-DA score plots (LV 1 vs. LV 2 and LV 1 vs. LV 3) illustrating the multivariate separation between samples collected at baseline (circles; “Pre-”) and following cocaine administration (squares; “Post-”). Experimental cohorts are color-coded by phenotype and sex: male bLRs (red), male bHRs (yellow), female bLRs (green), and female bHRs (blue). Shaded ellipses denote the 95 % confidence intervals for each group. **b)** Heatmap representing the fold-change of 230 metabolites identified with a significant main effect of cocaine administration (mixed-effects model with FDR correction, *q* < 0.1). These metabolites also exhibited variable importance in projection (VIP) scores > 1 and are shown in descending order. **c)** Pathway analysis of the metabolites that were significantly altered by cocaine administration.

A total of 230 metabolites showed a significant main effect of cocaine exposure after FDR correction (*q* < 0.1), further highlighting its dominant influence over the brain extracellular metabolome (Table S4). Fig. 6B shows the fold change between post- and pre-cocaine administration samples for each sex-phenotype combination. The metabolites shown in Fig. 6B are in descending order based on their VIP score from the PLS-DA model. Notably, all 230 significantly altered metabolites following cocaine administration exhibited VIP scores > 1, identifying them as key drivers of the separation in the PLS-DA model. Several compounds associated with cocaine metabolism were detected in brain dialysate and displayed significant sex-based differences (Fig. S6). These compounds were incorporated into the PLS-DA model (Fig. 6A) to illustrate global brain metabolome shifts, reflecting their high VIP scores within the model. Pathway enrichment analysis also revealed that cocaine administration broadly impacted amino acid metabolism (specifically arginine, alanine, aspartate, and glutamate pathways), purine metabolism, and arachidonic acid metabolism (Fig. 6C).

Building on the analysis in Fig. 6C, we first show how the individual compounds within these disrupted pathways shifted following cocaine exposure. Notably, cocaine administration impacted several pathways that originally differentiated the phenotypes at baseline (Fig. 5A-B), including arginine biosynthesis as well as alanine, aspartate, and glutamate metabolism (Fig. 7A). Mixed effects analysis found that glutamine only had a main effect of cocaine (*F*_1,28_ = 48.77, *p* < 0.0001). Meanwhile, three metabolites had main effects based on phenotype (see Fig. 5A-B for results), cocaine administration, and phenotype × administration interaction: arginine (cocaine: *F*_1,28_ = 66.62, *p* < 0.0001; interaction: *F*_1,28_ = 7.28, *p* = 0.01), *N*-acetylaspartate (cocaine: *F*_1,28_ = 146.7, *p* < 0.0001; interaction: *F*_1,28_ = 14.66, *p* = 0.0007), and NAAG (cocaine: *F*_1,28_ = 100.9, *p* < 0.0001; interaction: *F*_1,28_ = 20.55, *p* < 0.0001). Interestingly, post hoc analysis revealed no significant phenotypic differences for these compounds post-administration (Fig. S7), suggesting a differential impact of cocaine on these pathways between phenotypes. Purine metabolism, which encompasses a suite of compounds that support energetic demands, genetic coding, and inter- and intracellular signaling, was also affected by cocaine.^24,25^ Five compounds related to purine metabolism had main effects of cocaine administration (Fig. 7B): allantoic acid (*F*_1,28_ = 121.0, *p* < 0.0001), guanosine monophosphate (*F*_1,28_ = 108.9, *p* < 0.0001), adenosine monophosphate (*F*_1,28_ = 52.34, *p* < 0.0001), adenosine (*F*_1,28_ = 177.1, *p* < 0.0001), and inosine (*F*_1,28_ = 52.17, *p* < 0.0001). Levels for all these metabolites increased after cocaine administration except for inosine, which is a byproduct of adenosine degradation. Lastly, Fig. 7C demonstrates that cocaine administration also produced alterations to arachidonic acid metabolism, which regulates synaptic signaling and neurotransmission.^60^ Post-administration, decreased signals were observed for 20-hydroxyarachidonic acid (*F*_1,28_ = 57.82, *p* < 0.0001), 5-oxo-eicosatetraenoic acid (5-oxoETE; *F*_1,28_ = 29.36, *p* < 0.0001), 2,3-dinor-8-iso prostaglandin F2α (2,3-dinor-8-iso PGF2α; *F*_1,28_ = 41.51, *p* < 0.0001), and 15-deoxy-Δ12,14-prostaglandin J2 (15d-PGJ2; *F*_1,28_ = 168.3, *p* < 0.0001).

**Figure 7.**
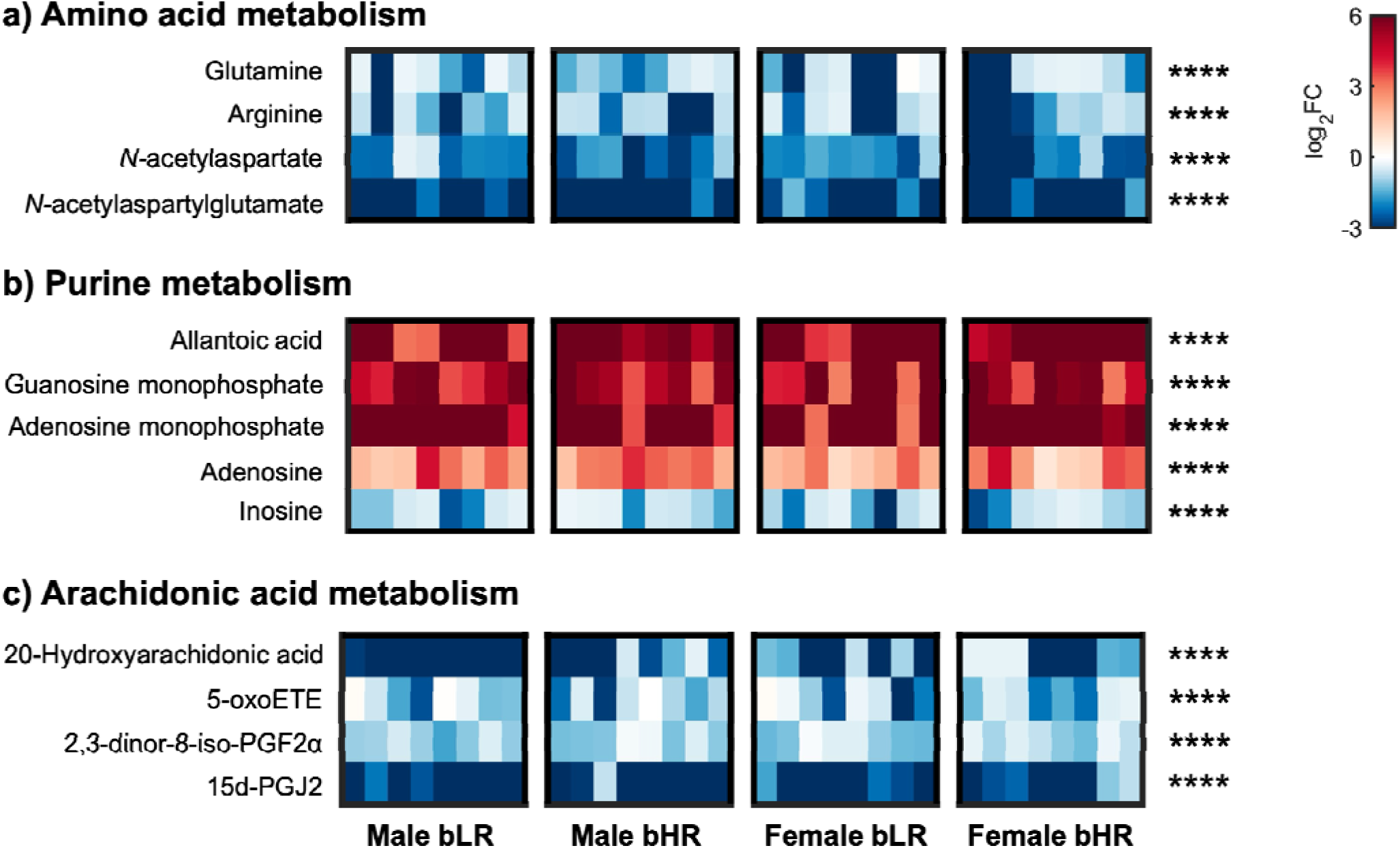
Primary metabolites associated with cocaine-induced shifts in biochemical pathways. Heatmaps showing the measured fold change (post-/pre-cocaine administration) across phenotype-sex groups. Each square within a phenotype-sex cohort represents an individual animal. Metabolites are categorized by their assigned pathway: **a)** amino acid metabolism; **b)** purine metabolism; and **c)** arachidonic acid metabolism. The color scale ranges from −3.0 to +6.0, where color represents the log_2_-fold change of the metabolite intensities. Asterisks indicate the main effect of cocaine administration from a three-way mixed effects model: **** *p* < 0.0001.

Beyond primary metabolic pathways, cocaine administration significantly altered the broader neurochemical landscape, specifically impacting biomarkers associated with cognitive function. Lipids serve as key cellular signaling molecules^26,61,62^ and several of these compounds demonstrated a robust main effect of cocaine administration. For example, Fig. 8A demonstrates that levels for 2-arachidonoyl glycerol ether (*F*_1,28_ = 48.36, *p* < 0.0001), docosanamide (*F*_1,28_ = 82.11, *p* < 0.0001), and 4-oxoretinoic acid (*F*_1,28_ = 155.6, *p* < 0.0001) all decreased post-administration. 4-oxoretinoic acid also exhibited significant sex × administration (*F*_1,28_ = 15.57, *p* = 0.0005) and phenotype × administration (*F*_1,28_ = 6.91, *p* = 0.01) interactions. Post hoc comparisons revealed that the decrease in metabolite levels was most pronounced in male bLR animals (*p* < 0.0001). Similarly, consistent with trends observed in Fig. 7A, several other amino acids and peptides with neuroprotective activity^63–65^ decreased post-administration. Exemplary compounds include pyroglutamic acid (*F*_1,28_ = 51.18, *p* < 0.0001), *N-*acetylglutamine (*F*_1,28_ = 101.0, *p* < 0.0001), and carnosine (*F*_1,28_ = 35.64, *p* < 0.0001; Fig. 8B). Lastly, Fig. 8C shows the signals measured for three additional metabolites that were categorized into different ClassyFire compound classes than those above. Two compounds that support cognitive function, *N*-acetylneuraminic acid^66^ (*F*_1,28_ = 53.47, *p* < 0.0001) and niacinamide^67^ (*F*_1,28_ = 51.18, *p* < 0.0001), decreased after cocaine administration. Meanwhile, hippuric acid, a byproduct of cocaine detoxification and aging,^68,69^ was found to increase post-exposure (*F*_1,28_ = 74.44, *p* < 0.0001).

**Figure 8.**
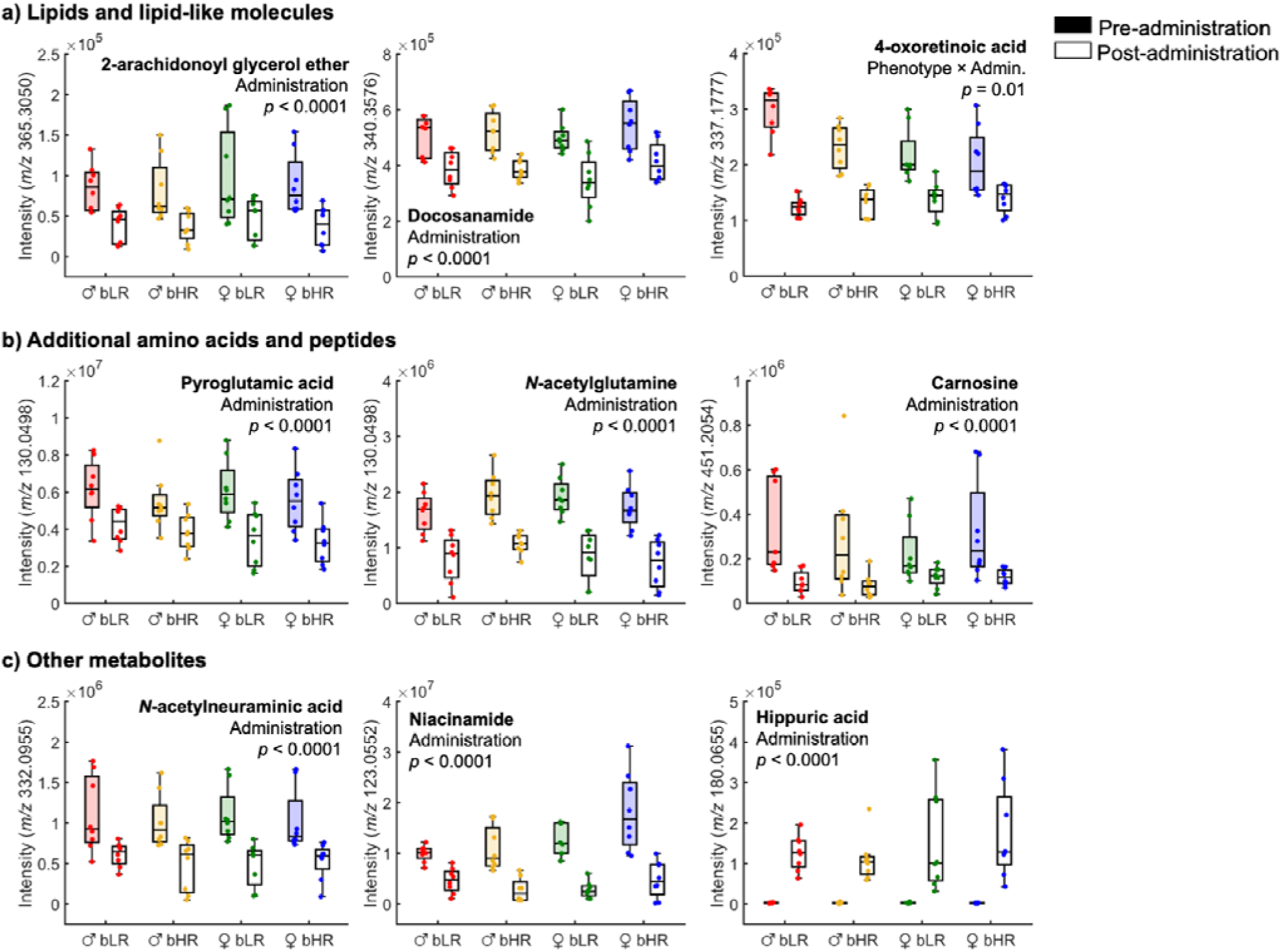
Impact of cocaine on metabolites associated with cognitive function. Box plots of the relative intensity measured in each experimental cohort for compounds related to **a)** lipid signaling, **b)** neuroprotective amino acids and peptides, and **c)** other exemplary metabolites. Box plots are shown with center line as median, extending to the IQR, with whiskers to the highest and lowest non-outlier points. Each dot represents data from a single animal (*N* = 8 rats/group). Calculated *p*-values for either the main effect of drug administration or phenotype × administration interaction is also provided.

## DISCUSSION

Analysis of the brain extracellular space can capture complex and dynamic neurochemical changes within a subject, reflecting the interplay of behavior, diseases, and pharmacology. We profiled the *in vivo* neurochemical landscape of the dorsomedial striatum and nucleus accumbens in bHR and bLR rodents, an established model of SUD vulnerability.^38^ Overall, we identified 680 compounds present in brain dialysate, expanding the extracellular metabolome from previous *in vivo* studies (Fig. 1).^28–34^ These identified metabolites span diverse chemical classes, ranging from small amino acids to larger lipid species. Hence, our untargeted analysis of brain dialysate captures a comparable chemical diversity to *ex vivo* benchmarks of the brain metabolome.^15^ While previous metabolomic studies of personality traits relied on *ex vivo* tissue or blood samples,^70–72^ this work is the first untargeted study to map brain neurochemistry dynamics *in vivo* across distinct behavioral phenotypes. Our metabolomic profiling readily differentiated these temperament phenotypes (Fig. 2), which was driven by metabolites mapping to pathways governing stress response, bioenergetics, and amino acid metabolism (Figs. 3-5). Acute cocaine administration reshapes regional brain chemistry, driving significant alterations in 230 extracellular metabolites (Fig. 6). Compared to previous studies on SUDs,^22–26,49^ our work offers the most comprehensive analysis of how cocaine impacts the brain metabolome. Functioning as a neurochemical “equalizer,” acute cocaine exposure masked many of the baseline phenotypic differences in amino acid metabolism, highlighting a differential effect on the bHR/bLR brain metabolome (Fig. 7). Cocaine administration also produced phenotype-independent alterations in purine processing, arachidonic acid pathways, and neuroprotective metabolites (Figs. 7-8). By capturing a broader array of localized signaling events than previous *ex vivo* approaches, this work underscores the necessity of *in vivo* monitoring to uncover mechanisms underlying SUD and other brain-related diseases.

While the bHR/bLR lines were selectively-bred for differences in novelty-induced locomotion, this model also shows concordant differences in stress response.^37,42^ Previous work has shown that hippocampal glucocorticoid receptors mRNA expression is higher in bLRs,^37,42^ contributing to increased anxiety-like behaviors.^73^ Yet basal plasma corticosterone levels are not statistically different between bHRs and bLRs,^42,74^ suggesting that peripheral measurements may not reflect neurochemical activity. Our *in vivo* measurements of the dorsomedial striatum and nucleus accumbens demonstrate that extracellular levels of corticosterone are elevated in bLRs (Fig. 3). We also identified phenotype-based associations with two other metabolites responsible for regulating stress response: LPA 18:1 and arachidonoyl ethanolamide (Fig. 3). LPA 18:1 was present in dialysate from bLRs but not detectable in bHRs. LPAs have been identified as key regulators in neuronal function, synaptic plasticity, and emotional regulation.^53–55^ Interestingly, intracranial administration of LPA 18:1 was found to increase anxiety-like behaviors in novel environments,^53^ matching observations of the bLR phenotype, suggesting that this lipid may be important in the behavioral differences observed. The endocannabinoid system also regulates emotional responses. In dialysate, not only did we observe detectable endocannabinoid content,^75^ but we found that arachidonoyl ethanolamide was significantly decreased in bLRs. Prior findings showed that a decline in arachidonoyl ethanolamide is also correlated with anxiety-like behaviors.^56,57^ Notably, alterations in both the LPA and endocannabinoid systems have not been previously implicated in bHR/bLR phenotypes. Hence, these findings open novel avenues for investigating signaling mechanisms underlying individual differences in stress-related molecules.

Within the studied reward-processing regions, we also found that the bHR and bLR animals exhibit basal extracellular differences in bioenergetic metabolites (Fig. 4). Acylcarnitines, essential transporters for mitochondrial fatty acid oxidation,^58^ and TCA cycle intermediates were significantly lower in bLRs. Meanwhile, bHRs exhibited elevated extracellular levels of metabolites characteristic of incomplete fatty acid oxidation.^59^ Previous studies have demonstrated that disruptions in energy production are correlated with internalizing disorders.^58,70,76^ Multi-omics evaluation of murine anxiety models reveals that shifting metabolic reliance from glycolysis to oxidative phosphorylation accompanies behavioral alterations.^70^ Within our rodent model, bLRs upregulate pathways driving fatty acid oxidation and TCA cycle activity, whereas bHRs maintain mitochondrial homeostasis by strictly regulating energy production.^48^ These distinct transcriptomic profiles directly relate to our metabolomic findings: emphasis on energy production in bLRs corresponds to lower acylcarnitine and TCA intermediate pools, while tighter gene control in bHRs yields elevated extracellular levels of bioenergetic metabolites and incomplete fatty acid oxidation products. Although direct intracellular measurements are ultimately required to validate these metabolic shifts, our study provides compelling evidence of a dynamic interplay between underlying genetic differences and metabolomic variations in bioenergetics.

Disruptions in brain mitochondrial function can also alter stress response and neurotransmission. Regarding the latter, mitochondria govern neurotransmitter dynamics by supplying key metabolic substrates and hosting enzymes for transmitter synthesis and degradation.^76,77^ For example, outer mitochondrial membrane monoamine oxidases directly regulate monoamine availability and turnover.^77,78^ Phenotypic alterations in mitochondrial function, and thus monoamine regulation, offer an additional mechanistic basis for our previous observation of significantly higher extracellular dopamine and norepinephrine levels in bHRs.^49^ Beyond monoamines, our current study uncovers broader metabolic divergences in amino acid pathways between phenotypes. Specifically, several metabolites associated with alanine, aspartate, and glutamate metabolism; arginine biosynthesis; glycine, serine, and threonine metabolism; branched-chain amino acids (valine, leucine, and isoleucine); and tryptophan metabolism were significantly elevated in bHRs (Fig. 5). These amino acid pathways serve as a direct link between baseline bioenergetics to neurotransmission, which can influence anxiety-like behavior.^71,72,77,79^ Collectively, the baseline extracellular differences in Figs. 3-5 suggest that mitochondrial dysfunction forces bLRs to consume energy at a disproportionate rate to regulate stress responses and maintain neurotransmission.

Notably, acute cocaine exposure exerted an “equalizing” effect on arginine biosynthesis and alanine, aspartate, and glutamate metabolism, driving extracellular levels of these metabolites down to uniform levels across the phenotypes (Fig. 7A). While similar alterations in excitatory amino acid pathways have been reported in *ex vivo* models,^23,25^ those studies examined cocaine effects broadly without accounting for underlying differences in SUD vulnerability. This “equalizing” effect between behavioral phenotypes for glutamate metabolism is biologically significant because increases in extracellular glutamate drive cocaine sensitization and drug-seeking behavior.^80^ Normally, NAAG acts as an endogenous mGluR3 agonist to inhibit glutamate release and suppress cocaine self-administration.^81^ However, despite higher baseline levels in bHRs, acute cocaine exposure abruptly depleted extracellular NAAG across all animals, suggesting that cocaine differentially modulates this metabolite and others related to amino acid pathways. This rapid loss removes a key inhibitory brake on glutamate transmission, potentially contributing to longer-term neuroadaptations to cocaine. Whether this acute “equalizing” effect endures over time or gives way to the pre-existing metabolic differences remains an important question for future research.

Pathway enrichment analysis also highlighted cocaine-induced alterations in purine metabolism. Purines are essential to brain function, regulating energy metabolism, cellular growth, neurotransmission, and neuromodulation.^24,25^ While cocaine is well known to increase extracellular adenosine and subsequently modulate dopaminergic signaling,^82^ our findings expand on this by revealing a broader activation of the purinergic system. We found that extracellular levels of allantoic acid, guanosine monophosphate, adenosine monophosphate, and adenosine were significantly elevated after cocaine administration (Fig. 7B). Upregulation of these purine metabolites in the brain tissue has also been observed in studies after acute cocaine exposure and self-administration.^24,25^ Interestingly, our study observed a significant decrease in extracellular inosine (Fig. 7B). Adenosine is typically cleared from the extracellular space through its enzymatic conversion to inosine; however, cocaine may disrupt this process to drive the accumulation of adensoine.^83^

We also detected a marked reduction in extracellular metabolites involved in arachidonic acid processing (Fig. 7C), a central pathway regulating lipid-mediated signaling and dopamine modulation.^60^ While previous studies in mice had showed that sphingolipid, glycerophospholipids, and glycolipid metabolisms were modified in brain tissue by cocaine administration,^26^ the specific impairment of arachidonic acid signaling represents a novel finding. Of interest, we found that the extracellular levels of 2-arachidonoyl glycerol ether decreased after acute cocaine exposure (Fig. 8A). In addition to detecting another key endocannabinoid in brain dialysate,^75^ these findings provide evidence that cocaine significantly alters central endocannabinoid tone. Prior studies show that cocaine mobilizes endocannabinoids within the nucleus accumbens to disinhibit dopamine neurons and enhance dopamine release, thereby reinforcing its rewarding actions.^84^ Nevertheless, future studies are needed to resolve the precise mechanisms linking cocaine exposure to disrupted arachidonic acid and endocannabinoid metabolism.

Growing evidence suggests that chronic cocaine use accelerates brain atrophy and cognitive decline.^85–87^ For example, magnetic resonance imaging data shows that individuals with cocaine dependence experience gray matter loss at twice the rate of healthy, age-matched controls.^86^ Positron emission tomography data have also revealed structural alterations in the prefrontal cortex and hippocampus of mice exposed to a single dose of cocaine.^87^ Motivated by this evidence, we found several neuroprotective and neurotoxic compounds that exhibited significant alterations post-cocaine administration (Fig. 8). We found that antioxidants such as carnosine^63^ and niacinamide^67^ decreased post-administration. Because substance abuse is known to deplete total antioxidant capacity, it disrupts vital defense mechanisms against reactive oxygen species.^88^ Consistent with these results, other neuroprotective compounds like docosanamide,^61^ 4-oxoretinoic acid,^62^ pyroglutamic acid,^64^ and *N*-acetylglutamine^65^ also decreased after cocaine exposure. Furthermore, post-cocaine decreases in extracellular *N-*acetylneuraminic acid, a key sialic acid supporting the brain’s gray matter architecture, directly points towards acute structural degradation in regions critical for cognitive function.^66^ In parallel with these reductions, we observed a post-administration increase in hippuric acid, a shared biomarker between cocaine-induced neurotoxicity^68^ and biological aging.^69^ Previously, an *ex vivo* study found age-related declines in niacinamide, *N*-acetylglutamine, and *N*-acetylneuraminic acid within the murine basal ganglia.^15^ In conjunction with these previous studies, our work suggests that acute cocaine administration can impact neuroprotective metabolites. Lower antioxidant levels combined with increases in markers of neurotoxicity signal that metabolic signatures resembling cellular stress or premature aging could be initiated after a single exposure to cocaine.

In summary, the brain extracellular space is a chemically rich compartment containing signaling molecules and metabolites that reflect brain cell activity. By analyzing samples from this space, we have defined metabolomic profiles that differentiate temperament phenotypes associated with a model of SUD vulnerability and the brain’s acute response to cocaine. These results demonstrate that the genetic underpinnings of emotional temperament are directly reflected in the brain extracellular metabolome. The analysis identified two stress-related signaling lipids that have previously not been associated with these phenotypes, arachidonoyl ethanolamide and LPA 18:1, opening new avenues of research at the intersection of SUDs and emotional temperament. Acute cocaine administration revealed both widespread and divergent metabolomic responses in the brain across sex and phenotype. Disruptions in glutamate, purine, and arachidonic acid metabolism, alongside altered neuroprotective compound profiles, identify new molecular pathways modulated by cocaine that warrant further investigation for their involvement in cocaine use disorder or cocaine-induced brain aging. Together, this *in vivo* characterization of cocaine’s impact on the brain metabolome provides the groundwork for precision therapeutics and targeted interventions. By capturing neurochemical dynamics directly where communication occurs, *in vivo* measurements can be utilized to contextualize both existing *ex vivo* data sets as well as broader genomic, transcriptomic, and proteomic landscapes of other brain-related disorders.

## METHODS

### Subjects

All studies used bHR (*N =* 16) and bLR (*N =* 16) Sprague-Dawley rats from a selective breeding colony in the Akil lab (generations 77, 79, 81, and 83).^38^ Each behavioral phenotype was studied using equal numbers of male and female animals (*N* = 8 per sex in each phenotype). Animals within each generation were also scored for locomotor response to novelty to confirm behavioral phenotype using an open field test. Rats weighed between 228-450 g for this study. Animals were group-housed in a temperature- and humidity-controlled room using a reverse light/dark cycle (12 h/12 h; lights off at 6 AM) and they had free access to food and water. All animals were treated in accordance with the National Institutes of Health guidelines on an approved protocol by the University of Michigan’s Institutional Animal Care and Use Committee.

### Microdialysis probe implantation

Carprofen (5 mg/kg) and gentamicin (2 mg/kg) were administered to all animals subcutaneously prior to stereotaxic surgery. During surgery, all animals were anesthetized with isoflurane (induction: 5 %; maintenance: 1.5-2.5 %) and a body temperature of 37 °C was maintained using a heating pad. CMA 12 guide cannulas (Harvard Apparatus, Holliston, MA) were unilaterally implanted based on the following coordinates from bregma: +1.8 AP, +1.8 ML, and -4.0 DV. A second dose of carprofen was administered 24 h post-operative and all animals were allowed to recover for a minimum of three days. After recovery, CMA 12 Elite microdialysis probes (4 mm long membrane, 0.5 mm O.D., and 20 kDa cutoff) were inserted through the guide cannula 48 h prior to sample collection while animals were under isoflurane anesthesia.

### Dialysate collection and preparation

Experiments were carried out in darkness to coincide with the animals’ light cycle. On the day of sample collection, animals were placed into a Raturn bowl (BASi, West Lafayette, IN), with free access to food and water, and tethered to the microdialysis lines. Probes were flushed at 2 µL/min for 1 h with artificial cerebrospinal fluid (aCSF; 145 mM sodium chloride, 2.68 mM potassium chloride, 1.40 mM calcium chloride, 1.01 mM magnesium sulfate, 1.55 mM dibasic sodium phosphate, 0.45 mM monobasic sodium phosphate, and 0.25 mM ascorbic acid) using a syringe pump (Fusion 400, Chemyx, Stafford, TX). The flow rate was then reduced to 1 µL/min for another 1 hr prior to dialysate collection. Baseline (pre-cocaine administration) samples were collected on ice at 1 µL/min for 3 h. Afterwards, cocaine (15 mg/kg, i.p.) was administered and microdialysis sampling continued for an additional 3 h. All samples were then stored in a -80 °C freezer for later processing. Dialysate samples from each animal pre- and post-cocaine administration were analyzed with LC-MS/MS in their native concentration. A QC reference sample was constructed by pooling together equal fractions of all dialysate samples. To develop a MS/MS reference sample, an aliquot of the pooled dialysate samples was concentrated 10-fold using a nitrogen evaporator (BenchTop Lab Systems, St. Louis, MO).

### LC-MS/MS analysis

Chromatographic data was collected using a Thermo Fisher Scientific (Waltham, MA) Vanquish Horizon LC coupled to an Orbitrap ID-X MS using methods previously described.^34,35^ All samples were analyzed in a randomized order. RPLC separations used a Waters (Milford, MA) HSST3 C_18_ column (2.1 mm × 100 mm, 1.8 µm). Mobile phase A was water with 0.1 % v/v formic acid and mobile phase B was methanol with 0.025 % v/v formic acid. The following mobile phase gradient was used for RP-LC: 0-10 min, 0-99 % B; 10-17 min, 99 % B; 17-17.1 min, 0 % B; 17.1-20 min, 0 % B. The flow rate for the RPLC separations was 0.45 mL/min. HILIC separations used a Waters BEH Amide (2.1 mm × 100 mm, 1.7 µm) column. Mobile phase A consisted of 95 % 10 mM ammonium formate in water/5 % acetonitrile with 0.125 % v/v formic acid while mobile phase B was 5 % 10 mM ammonium formate in water/95 % acetonitrile with 0.125 % v/v formic acid. The HILIC mobile phase gradient program was: 0-0.5 min, 100 % B; 0.5-7 min, 85 % B; 7-9 min, 85 % B; 9-16 min, 50 % B; 16-16.1 min, 100 % B; 16.1-20 min, 100 % B. The flow rate for the HILIC separations was 0.3 mL/min. For all separations, the injection volume was 7 µL, the autosampler temperature was 4 °C, and the column temperature was 55 °C.

MS data was collected in either positive or negative ionization mode using spray voltages of +3200 V or -3200 V, respectively. Full scan MS data was collected using the following settings: sheath gas, 40; auxiliary gas, 10; sweep gas, 1; ion transfer tube temperature, 325 °C; vaporizer temperature, 300 °C; full scan Orbitrap resolution, 120000; scan range, 70-1100 *m/z*; RF lens, 45 %; normalized AGC target, 25 %; maximum injection time, 50 ms; microscans, 1; data type, profile; internal mass calibration, EASY-IC. Data-dependent MS/MS acquisition maintained the same instrument settings except for the full scan Orbitrap resolution. The MS/MS parameters were: full scan Orbitrap resolution, 60000; intensity threshold, 1×10^4^; dynamic exclusion, 3 s (after one occurrence in a 5 ppm window); isolation mode, quadrupole; isolation window, 1.2 *m/z*; activation type, HCD; collision energy mode, assisted; collision energies, 20, 40, and 80 %; detector type, Orbitrap; Orbitrap resolution, 30000; normalized AGC target, 20%; maximum injection time, 54 ms; microscans, 1; data type, centroid; cycle time, 1.2 s. Five iterative injections (i.e., rolling precursor ion exclusion) were performed to improve the collection of MS/MS spectra for low abundance and/or overlapped analytes.

### Data processing and statistics

Chromatograms were converted from the raw vendor format to .mzXML files using MSConvertGUI (ProteoWizard, Palo Alto, CA)^89^ and imported into Matlab 2024a (Mathworks, Inc., Natick, MA) for analysis. Full-scan LC-MS chromatograms were first processed using regions of interest (ROI) compression to obtain a two-way data structure and remove background noise. Chromatographic signals with a peak width ≥ 2 s, a mass deviation ≤ 5 ppm, and intensity ≥ 1×10^4^ were extracted and inserted into an empty chromatogram at the original retention times and average *m/z*. Afterwards, the chromatograms were baseline corrected using a rolling ball minimum and normalized to the sum of the total ion current chromatogram. Feature lists for each chromatographic data set were developed using tile-based F-ratio analysis, with a respective tile size of 12.5 s for the RPLC data and 25.5 s for the HILIC data. *LOF* clustering^35^ was then applied to those feature lists to group together degeneracies (e.g., isotopes, adducts, in-source fragments). A cluster window size equal to half the tile size and a *LOF* threshold ≤ 20 % was used. *LOF* clustering yielded an updated table for each chromatographic data set that consolidates all detected degeneracies into “feature groups”.

Compound identification for each “feature group” was performed by MS/MS matching using a 10× preconcentrated pooled dialysate sample. The observed full-scan *m/z* for features within each group were initially searched as a precursor ion in the MS/MS data collected using a 5 ppm tolerance. Experimental MS/MS spectra were then searched against public libraries, including NIST20 (National Institute for Standards and Technology, Gaithersburg, MD), LipidBlast, and MassBank of North America (MONA, www.mona.fiehnlab.ucdavis.edu). For MS/MS database searching, experimental and library precursor ions were required to be within 5 ppm of expected values. Match quality between experimental and library MS/MS spectra was quantified using both a dot product (DP) and spectral entropy score (ES). A DP > 700 and ES > 0.65 were required for reporting. When available, authentic standards were used to further confirm metabolite identifications, requiring a retention time difference ≤ 0.5 min. Hence, all identifications made herein are with Metabolomics Standards Initiative^51^ Level 1 or 2 confidence. Meanwhile, “feature groups” without a high-quality spectral match were labeled as unknowns. All identified “feature groups” were combined into a unified master list for final filtering. For metabolites detected across multiple platforms, the entry with the lowest RSD in the QC samples was retained. Additionally, metabolites detected in at least 80% of samples within any single class (defined by combinations of sex, phenotype, and drug administration) were kept for downstream analysis.

Statistical analyses were performed using the peak intensity from the precursor ion for each identified metabolite. Missing data was imputed by 1/5^th^ of the ROI signal threshold. To correct for batch effects and instrumental drift, metabolite intensities were corrected using systematic error removal by random forest.^90^ Prior to multivariate analysis, metabolomic data were log-transformed and auto-scaled. PLS-DA visualized sample clustering while PERMANOVA assessed the statistical separation of the sample classes. Homogeneity of multivariate dispersions was tested prior to PERMANOVA using the *betadisper* function in the *vegan* package of R (version 4.5.1), which confirmed that within-class metabolomic variance was homogenous across sample classes (*p* > 0.05). PERMANOVA was performed on Euclidean distances with 999 permutations using the *adonis2* (*vegan* package; R 4.5.1). To evaluate baseline metabolomic profiles, the PERMANOVA model specified main effects for sex and phenotype along with their interaction. For comparing pre-versus post-cocaine administration, the PERMANOVA model was expanded to include sex, phenotype, drug administration, and all full interaction terms, with permutations restricted within subject to account for the repeated-measures design.

Individual baseline metabolite differences associated with sex and/or phenotype were identified using a two-way ANOVA. To control for multiple hypothesis testing across all evaluated metabolites, *p-*values were false discovery rate (FDR) corrected using the Benjamini–Hochberg procedure. Compounds meeting an FDR-adjusted threshold of *q* < 0.1 for either sex or phenotype were compiled into their respective significant metabolite lists (Table S1-S2). Similarly, pre-vs. post-cocaine metabolomic changes were evaluated using a three-way mixed effects model (incorporating sex, phenotype, cocaine administration, and all full interaction terms as fixed effects, with individual animals modeled as a random effect). Metabolites altered by cocaine administration (Table S4) were identified by applying FDR correction (*q* < 0.1) to the main effect of cocaine from the mixed-effects model. Unadjusted *p-*values for these selected compounds are provided throughout the text. Using these selected compounds, pathway analysis was performed using MetaboAnalyst 6.0 (www.metaboanalyst.ca) using hypergeometric testing, relative-betweeness centrality, and the KEGG *Rattus norvegicus* library.

## DATA AVAILABILITY

The LC-MS data sets generated in this study are publicly available as open-source .mzXML files via Zenodo: https://doi.org/10.5281/zenodo.21796722. Due to substantial file sizes, vendor format Thermo .raw files are available upon request.

## CODE AVAILABILITY

All scripts and functions used to read the .mzXML files and perform subsequent analyses can be accessed in the same Zenodo repository: https://doi.org/10.5281/zenodo.21796722.

## ACKNOWLEDGEMENTS

This work was funded by the National Institutes of Health under the award numbers: F32DA061554 (C.N.C.), U01DA043098 (H.A.), and RF1NS128522 (R.T.K.). Additional support also comes from the Hope for Depression Research Foundation and Pritzker Neuropsychiatric Research Consortium (H.A.). LC-MS/MS resources were provided by the Biomedical Research Core Facilities (Metabolomics Core) at the University of Michigan.

## AUTHOR CONTRIBUTIONS

C.N.C., P.P., H.A., and R.T.K. conceptualized this project. E.K.H-B. and H.A. managed the bHR/bLR rat colony and provided the animals. C.R.E. provided access to the LC-MS/MS and its associated resources. C.N.C., P.P., and N.M.O. performed the stereotaxic surgeries and managed *in vivo* dialysate collection. C.N.C. and P.P. acquired the untargeted LC-MS/MS metabolomic data. J.C.V.L. acquired the LC-MS/MS data on available authentic standards and confirmed metabolite identifications. C.N.C, P.P., H.A., and R.T.K. contributed to data analysis and interpretation. C.N.C. wrote an initial draft of the manuscript and all authors contributed to its editing.

