## Supplementary Information for "Deep *in vivo* brain metabolomics maps vulnerability-associated and cocaine-induced neurochemical states"

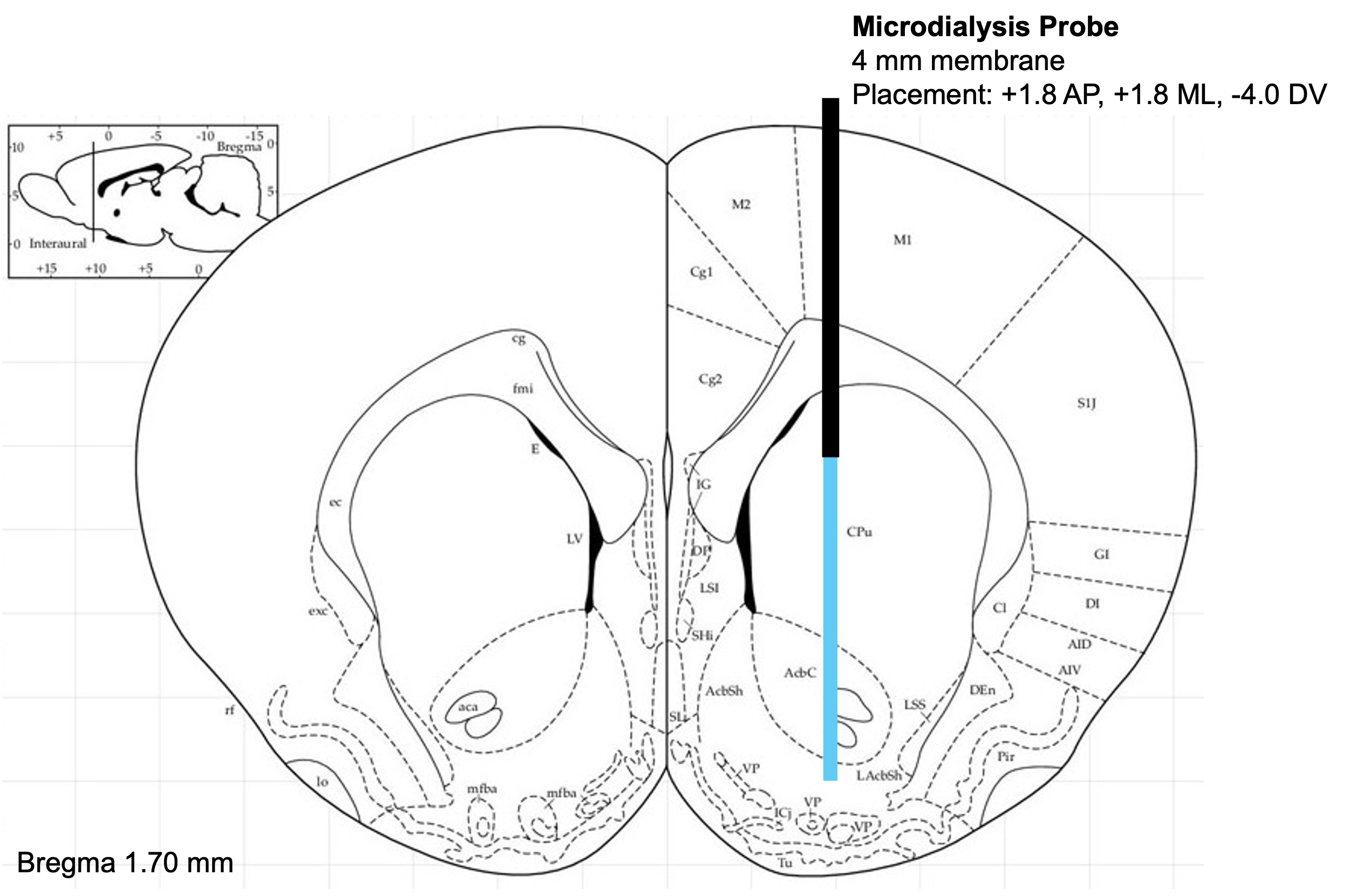

**Figure S1.** Illustration of the microdialysis probe placement. The guide cannula is represented by the black rectangle and the 4 mm microdialysis probe is represented by the blue rectangle. The stereotaxic coordinates are provided. Reference for the coronal rat brain slice was provided by Matt Gaidica (Washington University in St. Louis; https://labs.gaidi.ca/rat-brain-atlas/).

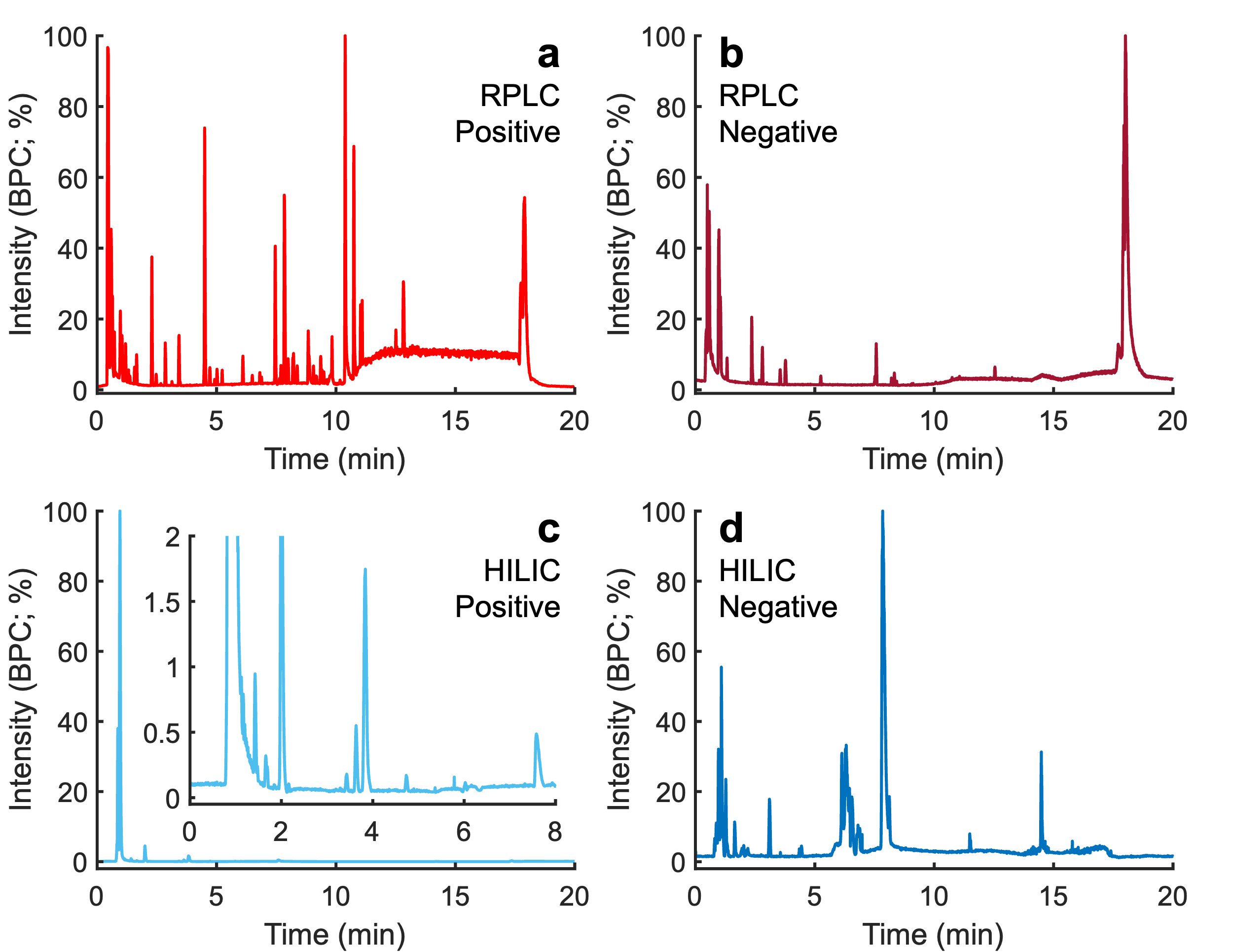

**Figure S2.** Representative base peak chromatograms (BPC) of brain dialysate in its native concentration. Data was collected using different combinations of chromatography and MS ionization modes: (A) reversed-phase liquid chromatography (RPLC) with positive ionization, (B) RPLC with negative ionization, (C) hydrophilic interaction liquid chromatography (HILIC) with positive ionization, and (D) HILIC with negative ionization.

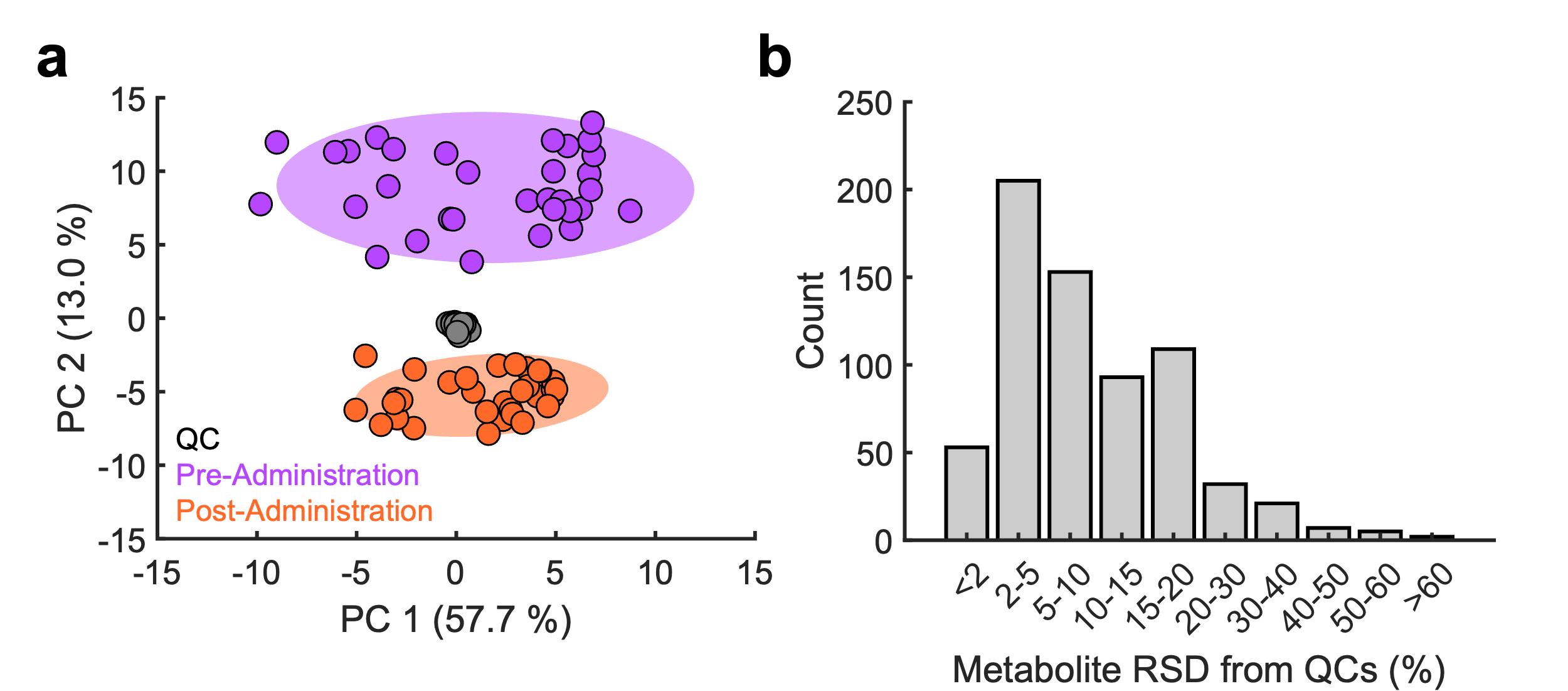

**Figure S3.** Quality control (QC) analysis for the dialysate metabolomic samples. (A) Principal components analysis (PCA) of the pre-administration (purple), post-administration (orange), and QC samples (gray). QC samples are highly clustered together, demonstrating low technical variance for this analysis. (B) Distribution of the technical variation of the annotated metabolites in the QC samples.

**Table S1. Compounds discovered to discriminate between males and females.** Dot product (DP) and entropy scores (ES) are provided as quality metrics for mass spectral library matching. An asterisk (*) indicates that these identifications were confirmed with standards.

| **Time (min)** | **Precursor *m/z*** | **Compound Identification** | **DP** | **ES** | **LC Mode** | **Ionization Mode** |
| --- | --- | --- | --- | --- | --- | --- |
| 0.59 | 203.0526 | 4-Amino-5-(2-phenylethyl)-4H-1,2,4-triazole-3-thiol | 994 | 0.81 | RPLC | POS |
| 0.62 | 114.0661 | * Creatinine | 1000 | 1.00 | RPLC | POS |
| 0.63 | 203.0155 | 4-(2-Thienyl)benzoic acid | 1000 | 1.00 | RPLC | POS |
| 0.63 | 265.1116 | * Thiamine cation | 911 | 0.98 | RPLC | POS |
| 0.73 | 140.0705 | 3-Amino-2,3-dihydrobenzoic acid | 789 | 0.76 | RPLC | POS |
| 0.73 | 175.1077 | N-alpha-Acetyl-L-ornithine | 824 | 0.77 | RPLC | POS |
| 0.87 | 155.0427 | 3-Ureidopropionic acid | 899 | 0.69 | RPLC | POS |
| 0.88 | 72.0807 | 1,2-Diamino-2-methylpropane | 962 | 0.99 | RPLC | POS |
| 0.92 | 84.0807 | 1,3-Dimethyl-6-(propylamino)-2,4(1H,3H)-pyrimidinedione | 940 | 0.68 | RPLC | POS |
| 0.94 | 204.123 | Acetyl-L-carnitine | 990 | 1.00 | RPLC | POS |
| 0.94 | 146.0923 | 4-Guanidinobutyric acid | 910 | 0.78 | RPLC | POS |
| 0.98 | 150.0549 | * Pyridoxal phosphate | 983 | 1.00 | RPLC | POS |
| 0.98 | 112.0504 | Cytidine 3'-monophosphate | 941 | 0.94 | RPLC | POS |
| 1.01 | 228.0978 | * 2'-Deoxycytidine | 1000 | 1.00 | RPLC | POS |
| 1.04 | 139.0501 | N-(3-Nitrophenyl)thiourea | 951 | 0.81 | RPLC | POS |
| 1.08 | 96.0443 | 4-Hydroxypyridine | 995 | 0.82 | RPLC | POS |
| 1.08 | 124.0392 | 4-Pyridinecarboxylic acid | 998 | 1.00 | RPLC | POS |
| 1.26 | 172.0402 | 4-Hydroxyquinoline-2-carbaldehyde | 943 | 0.70 | RPLC | POS |
| 1.27 | 126.0661 | * 5-Methylcytosine | 996 | 1.00 | RPLC | POS |
| 1.33 | 176.0293 | 4-Cyclohexyl-5-phenyl-2,4-dihydro-3H-1,2,4-triazole-3-thione | 938 | 0.68 | RPLC | POS |
| 1.35 | 87.044 | Crotonic acid | 1000 | 1.00 | RPLC | POS |
| 1.36 | 147.044 | 1,4-Pentadien-3-one, 1,5-bis(2-hydroxyphenyl)-, (1E,4E)- | 980 | 0.85 | RPLC | POS |
| 1.36 | 109.0647 | 6-Heptynoic acid | 842 | 0.82 | RPLC | POS |
| 1.36 | 123.044 | (-)-Catechin | 883 | 0.73 | RPLC | POS |
| 1.61 | 276.1442 | L-Glutarylcarnitine | 932 | 0.83 | RPLC | POS |
| 1.62 | 218.1386 | L-Propionylcarnitine | 995 | 1.00 | RPLC | POS |
| 1.62 | 132.1018 | * Isoleucine | 995 | 0.82 | RPLC | POS |
| 1.66 | 185.042 | 4-alpha-Mannobiose | 1000 | 1.00 | RPLC | POS |
| 1.76 | 108.0443 | 5-Hydroxy-2-(hydroxymethyl)pyridine | 988 | 0.93 | RPLC | POS |
| 1.77 | 153.0658 | 4-Methoxypyridine-2-carboxamide | 977 | 0.86 | RPLC | POS |
| 1.80 | 140.0342 | 6-Hydroxynicotinic acid | 929 | 0.76 | RPLC | POS |
| 1.81 | 168.0631 | (Diethylamino)(oxo)acetic acid | 989 | 0.80 | RPLC | POS |
| 2.43 | 232.1543 | Isobutyryl-L-carnitine | 999 | 1.00 | RPLC | POS |
| 2.75 | 100.0756 | N-[3-[Acetyl(hydroxy)amino]propyl]acetamide | 993 | 0.96 | RPLC | POS |
| 2.84 | 118.0651 | Indole | 892 | 0.65 | RPLC | POS |
| 3.04 | 138.0549 | 5-Methylpyridine-2-carboxylic acid | 908 | 0.78 | RPLC | POS |
| 3.14 | 246.0736 | Asn-Asp | 995 | 0.82 | RPLC | POS |
| 3.14 | 182.0811 | L-Tyrosine | 944 | 0.89 | RPLC | POS |
| 3.16 | 224.0917 | N-Acetyl-L-tyrosine | 952 | 0.89 | RPLC | POS |
| 3.18 | 246.1699 | Isovaleryl-L-carnitine | 996 | 1.00 | RPLC | POS |
| 3.19 | 144.0654 | Acetyl-L-threonine | 975 | 0.99 | RPLC | POS |
| 3.54 | 190.0499 | * Kynurenic acid | 944 | 0.81 | RPLC | POS |
| 3.58 | 295.1289 | Phe-Glu | 950 | 0.83 | RPLC | POS |
| 3.89 | 181.0835 | N5,N5-dimethyl[1,2,5]oxadiazolo[3,4-b]pyrazine-5,6-diamine | 790 | 0.82 | RPLC | POS |
| 4.03 | 127.0753 | 3,4-Dimethyl-1,2-cyclopentadione | 831 | 0.76 | RPLC | POS |
| 4.22 | 260.1855 | Hexanoyl-L-carnitine | 961 | 0.74 | RPLC | POS |
| 4.41 | 377.1458 | * (-)-Riboflavin | 987 | 0.90 | RPLC | POS |
| 4.81 | 208.0968 | N-Acetyl-L-phenylalanine | 992 | 0.88 | RPLC | POS |
| 4.82 | 146.06 | Resedine | 999 | 1.00 | RPLC | POS |
| 4.92 | 225.1097 | tert-Butyl 4-(6-amino-5-nitropyrimidin-4-yl)piperazine-1-carboxylate | 721 | 0.91 | RPLC | POS |
| 4.99 | 150.0913 | N-Cyclohexyl-3-phenylpropanamide | 929 | 0.78 | RPLC | POS |
| 5.03 | 131.0702 | 5-(1-Hydroxyethyl)oxolan-2-one | 903 | 0.81 | RPLC | POS |
| 5.25 | 283.1152 | 2-Aminoadenosine | 995 | 0.82 | RPLC | POS |
| 5.27 | 251.0525 | Met-Cys | 915 | 0.66 | RPLC | POS |
| 5.77 | 167.1066 | cis-Pinonic acid | 866 | 0.79 | RPLC | POS |
| 5.77 | 143.1066 | Cyclohexylacetic acid | 936 | 0.89 | RPLC | POS |
| 5.77 | 207.0991 | 7-Methoxy-6-methyl-2,4-pteridinediamine | 995 | 0.82 | RPLC | POS |
| 5.77 | 83.0855 | 2-Hexen-1-ol, (Z)- | 1000 | 1.00 | RPLC | POS |
| 5.91 | 289.2203 | L-Octanoylcarnitine | 906 | 0.93 | RPLC | POS |
| 5.95 | 319.1267 | N1-[3-(Benzyloxy)benzyl]-1H-tetrazole-1,5-diamine | 933 | 0.72 | RPLC | POS |
| 6.04 | 207.138 | 3-Methoxy-5-pentyl-2-prenylphenol | 995 | 0.82 | RPLC | POS |
| 6.89 | 181.1223 | 2(3H)-Furanone, dihydro-3-(1-hydroxyhexyl)-4-(hydroxymethyl)- | 955 | 0.90 | RPLC | POS |
| 6.96 | 321.1017 | N4-Acetylsulfamethazine | 992 | 0.81 | RPLC | POS |
| 7.00 | 315.178 | (3-Hydroxy-2,6,6-trimethylcyclohex-1-en-1-yl)methyl-beta-D-glucopyranoside | 928 | 0.76 | RPLC | POS |
| 7.06 | 163.0753 | 1-(Benzoyloxy)propan-2-yl benzoate | 959 | 0.88 | RPLC | POS |
| 7.12 | 316.2485 | Decanoyl-L-carnitine | 1000 | 1.00 | RPLC | POS |
| 7.13 | 95.0854 | 1,2,3,6-Tetrahydrobenzylalcohol | 984 | 1.00 | RPLC | POS |
| 7.13 | 197.1536 | 5-(6-Methyl-7-oxooctyl)furan-2(5H)-one | 920 | 0.89 | RPLC | POS |
| 7.20 | 216.1958 | Dodecanoic acid, 12-[[(cyclohexylamino)carbonyl]amino]- | 762 | 0.80 | RPLC | POS |
| 7.29 | 209.1536 | 2(5H)-Furanone, 5-(7-hydroxy-6-methyloctyl)- | 824 | 0.81 | RPLC | POS |
| 7.41 | 359.2044 | Pro-Ser-Arg | 995 | 0.82 | RPLC | POS |
| 7.44 | 279.0902 | Sulfamethazine | 988 | 0.72 | RPLC | POS |
| 7.54 | 343.173 | PyroGlu-Gly-Arg | 974 | 0.79 | RPLC | POS |
| 7.70 | 211.1692 | 4-(2,6,6-Trimethyl-3-oxocyclohex-1-en-1-yl)butan-2-yl-beta-D-glucopyranoside | 753 | 0.93 | RPLC | POS |
| 7.70 | 175.1481 | (3E)-4-(2,4,6-Trimethyl-3-cyclohexen-1-yl)-3-buten-2-one | 911 | 0.86 | RPLC | POS |
| 7.80 | 227.1042 | 9H-Purine-2,6-diamine, N6-(1R,2S,4S)-bicyclo[2.2.1]hept-2-yl-N2-phenyl-, rel- | 990 | 0.81 | RPLC | POS |
| 7.81 | 361.2357 | 7(S),17(S)-Dihydroxy-8(E),10(Z),13(Z),15(E),19(Z)-docosapentaenoic acid | 961 | 0.70 | RPLC | POS |
| 8.26 | 233.1536 | Naphtho[2,3-b]furan-2(4H)-one, 4a,5,6,7,8,8a,9,9a-octahydro-3,8a-dimethyl-5-methylene- | 965 | 0.92 | RPLC | POS |
| 8.26 | 273.146 | 7-Hydroxycostic acid | 980 | 0.72 | RPLC | POS |
| 8.36 | 241.1799 | Tetradecanedioic acid | 753 | 0.92 | RPLC | POS |
| 8.43 | 341.1954 | (-)-1,2:5,6-Di-O-cyclohexylidene-L-inositol | 988 | 0.80 | RPLC | POS |
| 8.51 | 347.1467 | (1S,5R,6S)-5-(Acetyloxy)-3-(hydroxymethyl)-4-oxo-6-(propan-2-yl)cyclohex-2-en-1-yl (2E)-2-methylbut-2-enoate | 966 | 0.66 | RPLC | POS |
| 9.04 | 301.1779 | Adrenosterone | 928 | 0.76 | RPLC | POS |
| 9.92 | 293.2089 | 9-Oxo-11-(3-pentyl-2-oxiranyl)-10E-undecenoic acid | 990 | 0.81 | RPLC | POS |
| 10.74 | 316.1661 | Ile-Trp | 981 | 0.72 | RPLC | POS |
| 0.57 | 265.1475 | Laurylsulfuric acid | 961 | 0.70 | RPLC | NEG |
| 0.94 | 218.1033 | Pantothenic acid | 997 | 1.00 | RPLC | NEG |
| 1.27 | 119.0351 | Acetic acid, 2,2'-[oxybis(2,1-ethanediyloxy)]bis- | 974 | 0.85 | RPLC | NEG |
| 1.33 | 125.0246 | Dimethyl 4-hydroxy-4-methyl-2-(3-nitrophenyl)-6-oxo-1,3-cyclohexanedicarboxylate | 1000 | 1.00 | RPLC | NEG |
| 1.50 | 195.051 | D-Gluconic acid | 991 | 0.95 | RPLC | NEG |
| 1.76 | 227.056 | 4-Nitro-N-(2-pyrimidinyl)benzamide | 940 | 0.76 | RPLC | NEG |
| 1.81 | 129.0195 | trans-Glutaconic acid | 1000 | 1.00 | RPLC | NEG |
| 1.81 | 113.0246 | Diethyl 2-[(2-hydroxyanilino)methylene]malonate | 1000 | 1.00 | RPLC | NEG |
| 2.24 | 373.0995 | N-Succinyl-5-aminoimidazole-4-carboxamide ribose | 922 | 0.88 | RPLC | NEG |
| 2.88 | 181.0508 | p-Hydroxyphenyllactic acid | 993 | 0.96 | RPLC | NEG |
| 3.01 | 161.0458 | beta-D-Glucopyranoside, 4-[(4-carboxy-3-hydroxy-3-methyl-1-oxobutoxy)methyl]phenyl | 965 | 0.73 | RPLC | NEG |
| 3.05 | 283.068 | Xanthosine | 890 | 0.84 | RPLC | NEG |
| 3.09 | 176.0388 | N-Formyl-L-methionine | 863 | 0.68 | RPLC | NEG |
| 3.58 | 212.0023 | Indoxyl sulfate | 989 | 0.80 | RPLC | NEG |
| 3.83 | 217.0423 | 2-(Diphenylphosphoryl)-1,4-benzenediol | 867 | 0.73 | RPLC | NEG |
| 4.01 | 188.0566 | N-Acetyl-L-glutamic acid | 996 | 1.00 | RPLC | NEG |
| 5.90 | 163.0403 | Cimicifugic acid K | 1000 | 1.00 | RPLC | NEG |
| 1.43 | 126.0661 | * 1-Methylcytosine | 886 | 0.72 | HILIC | POS |
| 1.80 | 110.06 | 1-Phenyl-1,3,8-triazaspiro[4.5]decan-4-one | 875 | 0.68 | HILIC | POS |
| 1.95 | 153.0658 | 2-Methyl-4-nitroaniline | 735 | 0.74 | HILIC | POS |
| 5.49 | 137.0709 | 1-Methylnicotinamide cation | 883 | 0.88 | HILIC | POS |

**Table S2. Compounds discovered to discriminate between bHRs and bLRs.** Dot product (DP) and entropy scores (ES) are provided as quality metrics for mass spectral library matching. An asterisk (*) indicates that these identifications were confirmed with standards.

| **Time (min)** | **Precursor *m/z*** | **Compound Identification** | **DP** | **ES** | **LC Mode** | **Ionization Mode** |
| --- | --- | --- | --- | --- | --- | --- |
| 0.58 | 175.1188 | * DL-Arginine | 995 | 0.88 | RPLC | POS |
| 0.62 | 114.0661 | * Creatinine | 1000 | 1.00 | RPLC | POS |
| 0.63 | 265.1116 | * Thiamine cation | 911 | 0.98 | RPLC | POS |
| 0.64 | 154.0585 | * Creatine | 1000 | 1.00 | RPLC | POS |
| 0.72 | 406.1315 | * N-Acetyl-D-lactosamine | 979 | 0.71 | RPLC | POS |
| 0.73 | 175.1077 | N-alpha-Acetyl-L-ornithine | 824 | 0.77 | RPLC | POS |
| 0.78 | 259.0931 | * 5-Methyluridine | 806 | 0.71 | RPLC | POS |
| 0.87 | 155.0427 | 3-Ureidopropionic acid | 899 | 0.69 | RPLC | POS |
| 0.92 | 189.1233 | N-alpha-Acetyl-L-lysine | 824 | 0.79 | RPLC | POS |
| 0.94 | 204.123 | Acetyl-L-carnitine | 990 | 1.00 | RPLC | POS |
| 0.96 | 145.0494 | trans-2-Butene-1,4-dicarboxylic acid | 920 | 0.84 | RPLC | POS |
| 0.97 | 198.0373 | * N-Acetyl-L-aspartic acid | 1000 | 1.00 | RPLC | POS |
| 0.97 | 203.139 | Ala-Ile | 873 | 0.73 | RPLC | POS |
| 0.98 | 168.0655 | Pyridoxal | 983 | 1.00 | RPLC | POS |
| 0.99 | 116.0341 | * L-Asparagine | 922 | 0.85 | RPLC | POS |
| 0.99 | 88.0392 | * L-Serine | 1000 | 1.00 | RPLC | POS |
| 1.08 | 96.0443 | 4-Hydroxypyridine | 995 | 0.82 | RPLC | POS |
| 1.27 | 166.0723 | * 7-Methylguanosine | 919 | 0.83 | RPLC | POS |
| 1.36 | 165.0546 | L-Tyrosine tert-butyl ester | 945 | 0.83 | RPLC | POS |
| 1.52 | 327.0792 | * Ac-Asp-Glu | 968 | 0.84 | RPLC | POS |
| 1.61 | 276.1442 | L-Glutarylcarnitine | 932 | 0.83 | RPLC | POS |
| 1.62 | 218.1386 | L-Propionylcarnitine | 995 | 1.00 | RPLC | POS |
| 1.62 | 132.1018 | * Isoleucine | 995 | 0.82 | RPLC | POS |
| 1.76 | 154.0498 | 3-Hydroxyanthranilic acid | 1000 | 1.00 | RPLC | POS |
| 1.77 | 151.0614 | Adonitol | 968 | 0.78 | RPLC | POS |
| 1.77 | 152.0566 | 1-{2-[(2-Amino-6-hydroxy-9H-purin-9-yl)methoxy]ethoxy}-3-methyl-1-oxo-2-butanamine | 955 | 0.87 | RPLC | POS |
| 2.43 | 232.1543 | Isobutyryl-L-carnitine | 999 | 1.00 | RPLC | POS |
| 2.57 | 279.1339 | Tyr-Pro | 778 | 0.66 | RPLC | POS |
| 2.84 | 130.065 | Indole-3-carbinol | 968 | 0.66 | RPLC | POS |
| 2.84 | 188.0706 | * L-Tryptophan | 999 | 1.00 | RPLC | POS |
| 2.84 | 146.06 | Indole-6-carboxaldehyde | 986 | 0.90 | RPLC | POS |
| 2.84 | 118.0651 | Indole | 892 | 0.65 | RPLC | POS |
| 3.18 | 246.1699 | Isovaleryl-L-carnitine | 996 | 1.00 | RPLC | POS |
| 3.25 | 229.1546 | Ile-Pro | 971 | 0.91 | RPLC | POS |
| 3.41 | 261.1446 | Leu-Glu | 968 | 0.72 | RPLC | POS |
| 4.22 | 260.1855 | Hexanoyl-L-carnitine | 961 | 0.74 | RPLC | POS |
| 5.27 | 251.0525 | Met-Cys | 915 | 0.66 | RPLC | POS |
| 5.91 | 289.2203 | L-Octanoylcarnitine | 906 | 0.93 | RPLC | POS |
| 7.04 | 177.091 | (2E,5E)-3,5,7-Trimethylocta-2,5-dienedioic acid | 827 | 0.76 | RPLC | POS |
| 7.06 | 163.0753 | 1-(Benzoyloxy)propan-2-yl benzoate | 959 | 0.88 | RPLC | POS |
| 7.12 | 316.2485 | Decanoyl-L-carnitine | 1000 | 1.00 | RPLC | POS |
| 7.41 | 359.2044 | Pro-Ser-Arg | 995 | 0.82 | RPLC | POS |
| 7.54 | 343.173 | PyroGlu-Gly-Arg | 974 | 0.79 | RPLC | POS |
| 7.61 | 347.2201 | * Corticosterone | 977 | 0.95 | RPLC | POS |
| 8.00 | 419.2556 | * 1-Oleoyl-L-alpha-lysophosphatidic acid | 969 | 0.65 | RPLC | POS |
| 8.36 | 241.1799 | Tetradecanedioic acid | 753 | 0.92 | RPLC | POS |
| 9.92 | 293.2089 | 9-Oxo-11-(3-pentyl-2-oxiranyl)-10E-undecenoic acid | 990 | 0.81 | RPLC | POS |
| 0.59 | 145.0144 | 2-Oxopentanedioic acid | 935 | 0.73 | RPLC | NEG |
| 0.93 | 173.0093 | cis-Aconitic acid | 994 | 1.00 | RPLC | NEG |
| 0.95 | 129.0195 | * L-2-Hydroxyglutaric acid | 998 | 1.00 | RPLC | NEG |
| 1.33 | 137.0246 | Hydroxysydonic acid | 960 | 0.73 | RPLC | NEG |
| 1.38 | 159.1028 | 3-Hydroxyoctanoic acid | 777 | 0.69 | RPLC | NEG |
| 1.70 | 115.0402 | 3-Oxopentanoic acid | 995 | 1.00 | RPLC | NEG |
| 2.70 | 131.0715 | * 2-Ethyl-2-hydroxybutyric acid | 1000 | 1.00 | RPLC | NEG |
| 3.09 | 176.0388 | N-Formyl-L-methionine | 863 | 0.68 | RPLC | NEG |
| 6.42 | 129.0559 | 3-Methyl-2-oxopentanoic acid | 950 | 0.77 | RPLC | NEG |
| 8.95 | 159.0301 | 2-Oxohexanedioic acid | 887 | 0.67 | RPLC | NEG |
| 0.88 | 179.143 | Dodeca-2(E),4(E)-dienoic acid | 902 | 0.87 | HILIC | POS |
| 0.93 | 348.2899 | * Arachidonoyl ethanolamide | 801 | 0.80 | HILIC | POS |
| 4.05 | 104.1069 | Glycerophosphocholine | 946 | 0.84 | HILIC | POS |
| 4.98 | 455.1889 | 2'-Deoxycytidine | 1000 | 1.00 | HILIC | POS |
| 4.98 | 112.0505 | Cytidine 3'-monophosphate | 924 | 0.83 | HILIC | POS |
| 6.86 | 144.1019 | Stachydrine | 993 | 1.00 | HILIC | POS |
| 7.04 | 118.0862 | Betaine | 1000 | 1.00 | HILIC | POS |

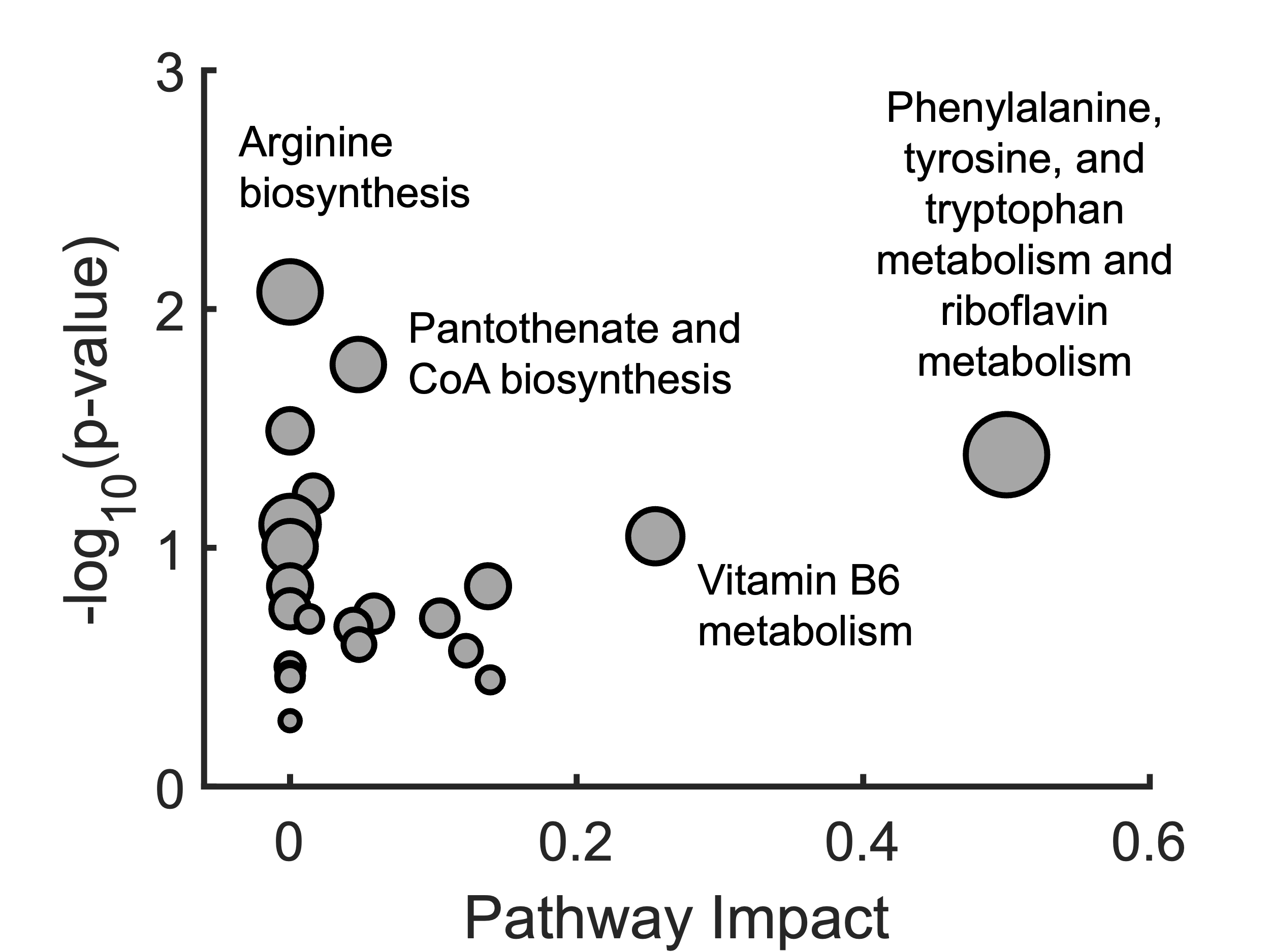

**Figure S4.** Pathway enrichment analysis of metabolites discovered in baseline dialysate samples with statistically significant sex-based differences. Note, the metrics for phenylalanine, tyrosine, and tryptophan metabolism and riboflavin metabolism are the same. These pathways highlight differences in neurotransmission and energy production between males and females.

**Table S3.** Two-way ANOVA results for detected acylcarnitines discovered in brain dialysate under baseline conditions.

| **Compound** | ***p-*value** | | |
| --- | --- | --- | --- |
|  | **Sex** | **Phenotype** | **Interaction** |
| Acetylcarnitine (C2) | 0.003 | 0.01 | 0.2 |
| Propionylcarnitine (C3) | 0.0008 | 0.0005 | 0.02 |
| Butyrylcarnitine (C4) | 0.009 | 0.03 | 0.04 |
| Isovalerylcarnitine (C5) | 0.01 | 0.009 | 0.08 |
| Hexanoylcarnitine (C6) | 0.002 | 0.01 | 0.08 |
| Decanoylcarnitine (C10) | 0.01 | 0.03 | 0.4 |
| Glutarylcarnitine (C5-DC) | 0.002 | <0.0001 | 0.2 |
| Adipoylcarnitine (C6-DC) | 0.003 | <0.0001 | 0.02 |

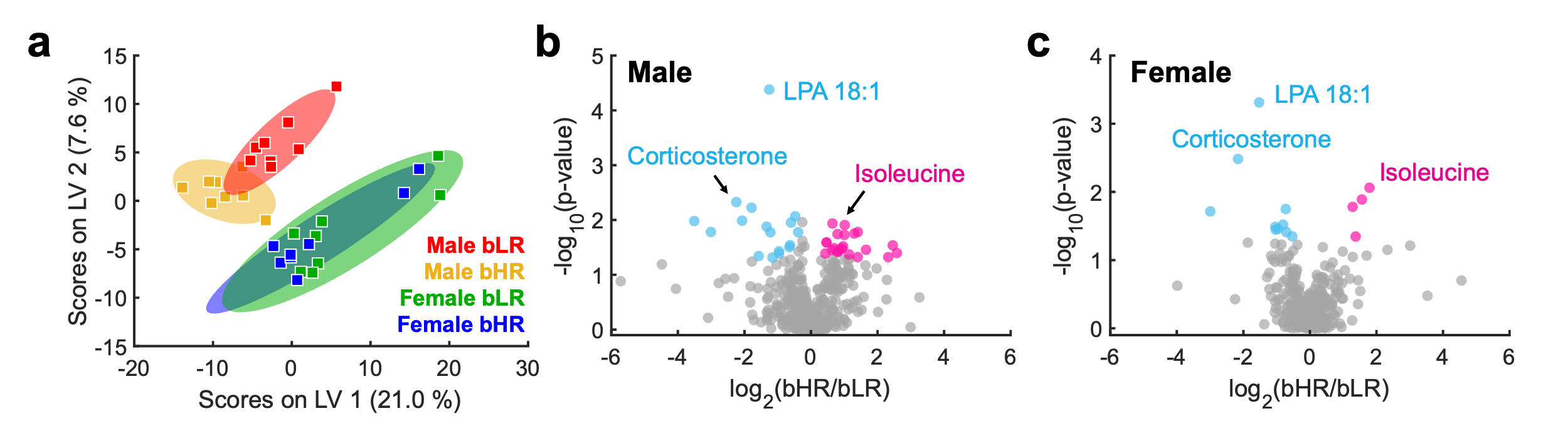

**Figure S5.** Evaluation of bHR/bLR phenotype differences in the dialysate samples acquired post-cocaine administration. (A) Partial least squares-discriminant analysis (PLS-DA) of the metabolomic samples. Experimental cohorts are color-coded by phenotype and sex: male bLRs (red), male bHRs (yellow), female bLRs (green), and female bHRs (blue). Shaded ellipses denote the 95 % confidence intervals for each group. (B) Volcano plot of metabolites discriminating male bLR and bHR rats post-cocaine administration. Gray points indicate non-significant features; teal and pink points denote metabolites enriched in bLRs and bHRs, respectively. Of the 64 phenotypic differences identified at baseline, 41 persisted following cocaine administration. (C) Volcano plot of metabolites discriminating female bLR and bHR rats post-cocaine administration. Of the 64 phenotypic differences identified at baseline, 15 persisted following cocaine administration. Across both sexes, compounds related to stress and bioenergetics primarily persisted after cocaine administration.
**Table S4. Compounds discovered to discriminate the dialysate samples before and after cocaine administration.** Dot product (DP) and entropy scores (ES) are provided as quality metrics for mass spectral library matching. An asterisk (*) indicates that these identifications were confirmed with standards.

| **Time (min)** | **Precursor *m/z*** | **Compound Identification** | **DP** | **ES** | **LC Mode** | **Ionization Mode** |
| --- | --- | --- | --- | --- | --- | --- |
| 0.56 | 175.1188 | * DL-Arginine | 995 | 0.88 | RPLC | POS |
| 0.57 | 130.0498 | D-Pyroglutamic acid | 1000 | 1.00 | RPLC | POS |
| 0.60 | 103.0389 | (2S)-4-Amino-2-hydroxybutyric acid | 920 | 0.82 | RPLC | POS |
| 0.62 | 332.0955 | * N-Acetylneuraminic acid | 984 | 0.86 | RPLC | POS |
| 0.63 | 217.0682 | (5-Ethylthiophen-2-yl)(phenyl)methanone | 984 | 0.67 | RPLC | POS |
| 0.63 | 203.0155 | 4-(2-Thienyl)benzoic acid | 1000 | 1.00 | RPLC | POS |
| 0.63 | 265.1116 | * Thiamine cation | 911 | 0.98 | RPLC | POS |
| 0.64 | 147.0763 | * L-Glutamine | 969 | 0.84 | RPLC | POS |
| 0.72 | 144.1018 | 1-Aminocyclohexanecarboxylic acid | 986 | 0.88 | RPLC | POS |
| 0.73 | 140.0705 | 3-Amino-2,3-dihydrobenzoic acid | 789 | 0.76 | RPLC | POS |
| 0.74 | 365.1058 | alpha,beta-Trehalose | 946 | 0.77 | RPLC | POS |
| 0.74 | 116.0705 | N-Benzyl-L-methionine methyl ester | 798 | 0.78 | RPLC | POS |
| 0.74 | 157.047 | 2,3-Dihydroxy-3-methylbutyric acid | 894 | 0.67 | RPLC | POS |
| 0.75 | 165.0132 | 2-Amino-4-hydroxybenzothiazole | 986 | 0.80 | RPLC | POS |
| 0.77 | 186.1125 | * p-Hydroxybenzoylecgonine | 863 | 0.79 | RPLC | POS |
| 0.78 | 259.0931 | * 5-Methyluridine | 806 | 0.71 | RPLC | POS |
| 0.80 | 200.1281 | * Methylecgonine | 999 | 1.00 | RPLC | POS |
| 0.87 | 155.0427 | 3-Ureidopropionic acid | 899 | 0.69 | RPLC | POS |
| 0.88 | 219.0975 | Ala-Glu | 833 | 0.74 | RPLC | POS |
| 0.88 | 72.0807 | 1,2-Diamino-2-methylpropane | 962 | 0.99 | RPLC | POS |
| 0.90 | 217.1294 | N-alpha-Acetyl-L-arginine | 988 | 0.80 | RPLC | POS |
| 0.92 | 130.0862 | 1-Acetylpiperidine-2-carboxylic acid | 1000 | 1.00 | RPLC | POS |
| 0.92 | 84.0807 | 1,3-Dimethyl-6-(propylamino)-2,4(1H,3H)-pyrimidinedione | 940 | 0.68 | RPLC | POS |
| 0.92 | 189.1233 | N-alpha-Acetyl-L-lysine | 824 | 0.79 | RPLC | POS |
| 0.93 | 129.0657 | 5-Methyl-5,6-Dihydrouracil | 970 | 0.87 | RPLC | POS |
| 0.94 | 111.0552 | 6-Amino-2-pyridone | 917 | 0.80 | RPLC | POS |
| 0.95 | 113.0344 | 3-(2-Hydroxy-4-oxo-1(4H)-pyrimidinyl)propanoic acid | 992 | 1.00 | RPLC | POS |
| 0.97 | 203.139 | Ala-Ile | 873 | 0.73 | RPLC | POS |
| 0.97 | 198.0373 | * N-Acetyl-L-aspartic acid | 1000 | 1.00 | RPLC | POS |
| 0.98 | 348.0707 | * Adenosine 5'-monophosphate | 986 | 0.86 | RPLC | POS |
| 0.98 | 168.0655 | * Pyridoxal | 998 | 1.00 | RPLC | POS |
| 1.00 | 99.0075 | Maleic anhydride | 1000 | 1.00 | RPLC | POS |
| 1.00 | 158.1175 | trans-4-(Aminomethyl)cyclohexanecarboxylic acid | 973 | 0.79 | RPLC | POS |
| 1.01 | 150.0582 | 2-(3-Carboxypropionylamino)-4-methylsulfanylbutyric acid | 997 | 1.00 | RPLC | POS |
| 1.01 | 133.0317 | N-(2,4-Dinitrophenyl)-L-methionine | 704 | 0.67 | RPLC | POS |
| 1.01 | 228.0978 | * 2'-Deoxycytidine | 1000 | 1.00 | RPLC | POS |
| 1.02 | 364.0656 | * Guanosine 5'-monophosphate | 995 | 0.88 | RPLC | POS |
| 1.03 | 330.0733 | 4',5-Dihydroxy-3',6,7-trimethoxyflavone | 973 | 0.79 | RPLC | POS |
| 1.25 | 123.0552 | * Niacinamide | 983 | 0.92 | RPLC | POS |
| 1.26 | 130.0498 | N-Acetyl-L-glutamine | 1000 | 1.00 | RPLC | POS |
| 1.27 | 258.1084 | * Benserazide | 966 | 0.65 | RPLC | POS |
| 1.27 | 126.0661 | * 5-Methylcytosine | 996 | 1.00 | RPLC | POS |
| 1.33 | 176.0293 | 4-Cyclohexyl-5-phenyl-2,4-dihydro-3H-1,2,4-triazole-3-thione | 938 | 0.68 | RPLC | POS |
| 1.34 | 202.1074 | Pantothenic acid | 993 | 0.81 | RPLC | POS |
| 1.36 | 91.0541 | Benzyl alcohol | 971 | 0.79 | RPLC | POS |
| 1.36 | 95.0491 | (2E,4E)-Hexa-2,4-dienoic acid | 964 | 0.85 | RPLC | POS |
| 1.36 | 136.0756 | Norphenylephrine | 982 | 0.92 | RPLC | POS |
| 1.36 | 137.0596 | 1,2,3,4-Tetrahydro-6,7-isoquinolinediol | 997 | 1.00 | RPLC | POS |
| 1.36 | 147.044 | 1,4-Pentadien-3-one, 1,5-bis(2-hydroxyphenyl)-, (1E,4E)- | 980 | 0.85 | RPLC | POS |
| 1.52 | 327.0792 | * Ac-Asp-Glu | 968 | 0.84 | RPLC | POS |
| 1.62 | 86.0963 | 1,5-Pentanediamine | 995 | 0.87 | RPLC | POS |
| 1.66 | 185.042 | 4-alpha-Mannobiose | 1000 | 1.00 | RPLC | POS |
| 1.73 | 136.0617 | 3-(6-Amino-9H-purin-9-yl)propanoic acid | 968 | 0.88 | RPLC | POS |
| 1.77 | 268.1039 | * Adenosine | 992 | 0.87 | RPLC | POS |
| 1.78 | 137.0457 | * Inosine | 1000 | 1.00 | RPLC | POS |
| 1.85 | 151.0753 | 4-Hydroxy-3-methoxyphenethyl alcohol | 903 | 0.78 | RPLC | POS |
| 2.02 | 155.0814 | Cyclo(glycylprolyl) | 984 | 0.90 | RPLC | POS |
| 2.19 | 113.0232 | 3-(Methoxycarbonyl)bicyclo[2.2.1]hept-5-ene-2-carboxylic acid | 995 | 1.00 | RPLC | POS |
| 2.30 | 166.0862 | * DL-Phenylalanine | 1000 | 1.00 | RPLC | POS |
| 2.30 | 103.0541 | 2-Methylisoquinolin-2-ium cation | 999 | 1.00 | RPLC | POS |
| 2.30 | 120.0807 | 1,2,3,4-Tetrahydroisoquinolin-5-amine | 968 | 0.87 | RPLC | POS |
| 2.30 | 131.049 | (Z)-N-(2-Hydroxyethyl)-3-phenylacrylamide | 1000 | 1.00 | RPLC | POS |
| 2.30 | 107.049 | 4-Hydroxy-3-(3-methyl-2-butenyl)benzaldehyde | 869 | 0.69 | RPLC | POS |
| 2.75 | 100.0756 | N-[3-[Acetyl(hydroxy)amino]propyl]acetamide | 993 | 0.96 | RPLC | POS |
| 2.88 | 149.0265 | Methionine sulfoxide | 948 | 0.81 | RPLC | POS |
| 3.09 | 156.1019 | 2,2-Bis(hydroxymethyl)-3-quinuclidinone | 956 | 0.83 | RPLC | POS |
| 3.18 | 148.0756 | Kynuramine | 947 | 0.71 | RPLC | POS |
| 3.25 | 132.0443 | Zonisamide | 904 | 0.73 | RPLC | POS |
| 3.31 | 320.1495 | * m-Hydroxycocaine | 905 | 0.92 | RPLC | POS |
| 3.41 | 261.1446 | Leu-Glu | 968 | 0.72 | RPLC | POS |
| 3.58 | 306.1337 | * m-Hydroxybenzoylecgonine | 976 | 0.83 | RPLC | POS |
| 3.58 | 180.0655 | * Hippuric acid | 967 | 0.83 | RPLC | POS |
| 3.98 | 137.0596 | 4-Hydroxy-3-methoxybenzenemethanol | 960 | 0.86 | RPLC | POS |
| 4.02 | 261.0845 | Felbamate | 986 | 0.80 | RPLC | POS |
| 4.04 | 120.0443 | 3-Pyridylacetic acid | 998 | 1.00 | RPLC | POS |
| 4.16 | 299.1102 | Triethyl citrate | 991 | 0.81 | RPLC | POS |
| 4.19 | 290.1387 | * Benzoylecgonine | 897 | 0.85 | RPLC | POS |
| 4.29 | 276.123 | * Benzoylnorecgonine | 982 | 0.92 | RPLC | POS |
| 4.29 | 304.1544 | * Cocaine | 960 | 0.98 | RPLC | POS |
| 4.29 | 304.1647 | Trp-Val | 904 | 0.73 | RPLC | POS |
| 4.50 | 343.1315 | Tyr-Tyr | 991 | 0.81 | RPLC | POS |
| 4.51 | 126.0912 | N-[6,6-Dimethyl-5-(1-methylisonipecotoyl)-1,4-dihydropyrrolo[3,4-c]pyrazol-3-yl]-3-methylbutyramide | 935 | 0.72 | RPLC | POS |
| 4.75 | 135.044 | 3-Methoxy-N-(4-methyl-1,3-thiazol-2-yl)benzamide | 851 | 0.84 | RPLC | POS |
| 4.82 | 146.06 | Resedine | 999 | 1.00 | RPLC | POS |
| 4.90 | 319.1734 | Gly-Ser-Arg | 991 | 0.81 | RPLC | POS |
| 4.91 | 211.1441 | Cyclo(leucylprolyl) | 974 | 0.96 | RPLC | POS |
| 4.97 | 423.1378 | Eprosartan | 984 | 0.80 | RPLC | POS |
| 5.03 | 131.0702 | 5-(1-Hydroxyethyl)oxolan-2-one | 903 | 0.81 | RPLC | POS |
| 5.11 | 179.1178 | 3-(Benzylamino)propanamide | 927 | 0.67 | RPLC | POS |
| 5.13 | 193.0608 | 5-Methoxy-1H-indazole-3-carboxylic acid | 745 | 0.66 | RPLC | POS |
| 5.18 | 242.1306 | threo-Dihydrobupropion | 920 | 0.67 | RPLC | POS |
| 5.18 | 245.1284 | 3-Benzylhexahydropyrrolo[1,2-a]pyrazine-1,4-dione | 809 | 0.74 | RPLC | POS |
| 5.24 | 105.0334 | 2-[(6-Ethoxy-1,3-benzothiazol-2-yl)thio]-1-phenylethanone | 851 | 0.67 | RPLC | POS |
| 5.25 | 283.1152 | 2-Aminoadenosine | 995 | 0.82 | RPLC | POS |
| 5.27 | 292.1179 | Corey lactone aldehyde benzoate | 782 | 0.80 | RPLC | POS |
| 5.27 | 251.0525 | Met-Cys | 915 | 0.66 | RPLC | POS |
| 5.32 | 136.0756 | 1-[4-(Dimethylamino)phenyl]ethan-1-ol | 928 | 0.95 | RPLC | POS |
| 5.38 | 140.1069 | rac-Ethyl (1R,2S)-2-aminocyclopentane-1-carboxylate | 769 | 0.70 | RPLC | POS |
| 5.59 | 209.1148 | 2-(1-Hydroxycyclohexyl)butanoic acid | 973 | 0.71 | RPLC | POS |
| 5.91 | 191.1042 | Dihydralazine | 1000 | 1.00 | RPLC | POS |
| 5.95 | 319.1267 | N1-[3-(Benzyloxy)benzyl]-1H-tetrazole-1,5-diamine | 933 | 0.72 | RPLC | POS |
| 6.02 | 389.1323 | Bicyclomycin benzoate | 957 | 0.77 | RPLC | POS |
| 6.02 | 390.1356 | 2-Acetamido-2-deoxy-3-O-(alpha-L-fucopyranosyl)-D-glucopyranose | 974 | 0.71 | RPLC | POS |
| 6.03 | 195.0992 | 4-Ethyl-5-methyl-2-(1H-tetrazol-5-yl)-1,2-dihydro-3H-pyrazol-3-one | 977 | 0.71 | RPLC | POS |
| 6.04 | 207.138 | 3-Methoxy-5-pentyl-2-prenylphenol | 995 | 0.82 | RPLC | POS |
| 6.06 | 451.2054 | * L-Carnosine | 982 | 0.79 | RPLC | POS |
| 6.06 | 163.0389 | Chorismic acid | 982 | 0.86 | RPLC | POS |
| 6.18 | 369.1999 | His-Gly-Arg | 994 | 0.81 | RPLC | POS |
| 6.28 | 385.1473 | 2-(2-((Hexopyranosyloxy)methyl)-3-methylcyclopent-2-en-1-yl)-3-hydroxypropanoic acid | 991 | 0.81 | RPLC | POS |
| 6.59 | 329.1937 | Gly-Pro-Arg | 1000 | 1.00 | RPLC | POS |
| 6.63 | 109.0647 | Ethyl 6-isopropyl-2-methyl-4-oxo-2-cyclohexene-1-carboxylate | 843 | 0.72 | RPLC | POS |
| 6.70 | 212.107 | 4-Methyl-N-phenylbenzamide | 955 | 0.75 | RPLC | POS |
| 6.81 | 331.2094 | Val-Gly-Arg | 991 | 0.81 | RPLC | POS |
| 6.87 | 202.1226 | 5-Methoxy-alpha-ethyltryptamine | 995 | 0.82 | RPLC | POS |
| 7.00 | 315.178 | (3-Hydroxy-2,6,6-trimethylcyclohex-1-en-1-yl)methyl-beta-D-glucopyranoside | 928 | 0.76 | RPLC | POS |
| 7.04 | 177.091 | (2E,5E)-3,5,7-Trimethylocta-2,5-dienedioic acid | 827 | 0.76 | RPLC | POS |
| 7.06 | 163.0753 | 1-(Benzoyloxy)propan-2-yl benzoate | 959 | 0.88 | RPLC | POS |
| 7.15 | 385.22 | His-Thr-Lys | 989 | 0.80 | RPLC | POS |
| 7.20 | 163.1481 | 4-Hydroxydodecanoic acid | 745 | 0.67 | RPLC | POS |
| 7.29 | 261.1096 | 1,3-Diaminopropane-N,N,N',N'-tetraacetic acid | 979 | 0.88 | RPLC | POS |
| 7.29 | 387.2355 | 2-Furanacetic acid, tetrahydro-5-(2-hydroxypropyl)-alpha-methyl-, 2-[5-(1-carboxyethyl)tetrahydro-2-furanyl]-1-methylethyl ester | 1000 | 1.00 | RPLC | POS |
| 7.38 | 192.1383 | Diethyltoluamide | 944 | 0.77 | RPLC | POS |
| 7.41 | 359.2044 | Pro-Ser-Arg | 995 | 0.82 | RPLC | POS |
| 7.42 | 241.1335 | N-Benzyl-N-methyl-N'-phenylurea | 941 | 0.78 | RPLC | POS |
| 7.49 | 135.0803 | 3,4-Dimethylbenzaldehyde | 965 | 0.80 | RPLC | POS |
| 7.49 | 389.2518 | Asn-Lys-Lys | 1000 | 1.00 | RPLC | POS |
| 7.54 | 265.1798 | Dinor-12-oxophytodienoic acid | 987 | 0.72 | RPLC | POS |
| 7.54 | 343.173 | PyroGlu-Gly-Arg | 974 | 0.79 | RPLC | POS |
| 7.60 | 259.1903 | Diethyl sebacate | 881 | 0.82 | RPLC | POS |
| 7.61 | 253.1409 | (1R,2S,5R)-2-Isopropyl-5-methylcyclohexyl 2,2-dihydroxyacetate | 995 | 0.82 | RPLC | POS |
| 7.61 | 369.2039 | 12-Methoxycarnosic acid | 988 | 0.80 | RPLC | POS |
| 7.70 | 151.148 | (2E)-3,4,7-Trimethylocta-2,6-dien-1-ol | 906 | 0.75 | RPLC | POS |
| 7.70 | 255.1591 | 3-Furancarboxylic acid, tetrahydro-4-methylene-2-octyl-5-oxo-, (2R,3S)-rel- | 770 | 0.85 | RPLC | POS |
| 7.74 | 179.1066 | 9-(Methoxycarbonyl)dec-9-enoic acid | 732 | 0.69 | RPLC | POS |
| 7.80 | 245.1146 | Glu-Val | 995 | 0.82 | RPLC | POS |
| 7.80 | 227.1042 | 9H-Purine-2,6-diamine, N6-(1R,2S,4S)-bicyclo[2.2.1]hept-2-yl-N2-phenyl-, rel- | 990 | 0.81 | RPLC | POS |
| 7.84 | 226.1226 | N-Benzyl-4-methylbenzamide | 968 | 0.86 | RPLC | POS |
| 7.84 | 267.1226 | 1-(1-Methylindol-5-yl)-3-(3-pyridyl)urea | 761 | 0.75 | RPLC | POS |
| 7.90 | 451.2306 | Glu-Phe-Arg | 994 | 0.81 | RPLC | POS |
| 7.90 | 127.0389 | 1,3,5-Benzenetriol | 765 | 0.67 | RPLC | POS |
| 7.92 | 383.2201 | 12-(Acetyloxy)pimara-7,15-dien-18-oic acid | 965 | 0.77 | RPLC | POS |
| 8.00 | 235.1303 | 1-(4-(7H-Purin-6-yl)morpholin-2-yl)methanamine | 1000 | 1.00 | RPLC | POS |
| 8.02 | 195.138 | 2(3H)-Furanone, dihydro-4-(hydroxymethyl)-3-(1-hydroxy-5-methylhexyl)- | 727 | 0.93 | RPLC | POS |
| 8.03 | 240.1382 | N,N-Dibenzylacetamide | 996 | 1.00 | RPLC | POS |
| 8.10 | 305.1723 | 4-Methoxy-6-pentyl-2-prop-1-en-2-yl-2,3-dihydro-1-benzofuran-7-carboxylic acid | 938 | 0.66 | RPLC | POS |
| 8.20 | 315.1568 | Acetylvalerenolic acid | 971 | 0.66 | RPLC | POS |
| 8.22 | 105.0698 | 3,4-Dimethylbenzene-1-sulfonic acid | 920 | 0.95 | RPLC | POS |
| 8.26 | 233.1536 | Naphtho[2,3-b]furan-2(4H)-one, 4a,5,6,7,8,8a,9,9a-octahydro-3,8a-dimethyl-5-methylene- | 965 | 0.92 | RPLC | POS |
| 8.26 | 273.146 | 7-Hydroxycostic acid | 980 | 0.72 | RPLC | POS |
| 8.32 | 263.1252 | 1,2-Dithiolane-3-pentanamide, N,N'-1,3-propanediylbis- | 979 | 0.79 | RPLC | POS |
| 8.36 | 241.1799 | Tetradecanedioic acid | 753 | 0.92 | RPLC | POS |
| 8.37 | 279.0991 | Phe-Asp | 1000 | 1.00 | RPLC | POS |
| 8.37 | 105.0334 | N-(5-Phenethyl-[1,3,4]thiadiazol-2-yl)benzamide | 955 | 0.99 | RPLC | POS |
| 8.38 | 225.091 | 1,3-Diphenylpropane-1,3-dione | 964 | 0.86 | RPLC | POS |
| 8.41 | 197.0961 | (S,S)-(-)-Hydrobenzoin | 937 | 0.95 | RPLC | POS |
| 8.43 | 341.1954 | (-)-1,2:5,6-Di-O-cyclohexylidene-L-inositol | 988 | 0.80 | RPLC | POS |
| 8.45 | 207.138 | 3-Hydroxy-2-octylpentanedioic acid | 934 | 0.85 | RPLC | POS |
| 8.48 | 313.2363 | (9Z,12E)-15,16-Dihydroxyoctadeca-9,12-dienoic acid | 959 | 0.65 | RPLC | POS |
| 8.51 | 347.1467 | (1S,5R,6S)-5-(Acetyloxy)-3-(hydroxymethyl)-4-oxo-6-(propan-2-yl)cyclohex-2-en-1-yl (2E)-2-methylbut-2-enoate | 966 | 0.66 | RPLC | POS |
| 8.70 | 249.146 | 9-[2-(4-Morpholinyl)ethyl]-9H-purin-6-ylamine | 994 | 0.81 | RPLC | POS |
| 8.76 | 211.0965 | 5-Hydroxy-2,2,6,6-tetramethyl-4-[3-methyl-1-[2,4,6-trihydroxy-3-(2-methylpropanoyl)phenyl]butyl]cyclohex-4-ene-1,3-dione | 994 | 0.81 | RPLC | POS |
| 8.81 | 325.1997 | 2,3-Dinor-8-isoprostaglandin-F2alpha | 893 | 0.71 | RPLC | POS |
| 8.81 | 197.1536 | Dodeca-2(E),4(E)-dienoic acid | 807 | 0.72 | RPLC | POS |
| 9.03 | 151.1117 | trans-4,5-Epoxy-2(E)-decenal | 867 | 0.79 | RPLC | POS |
| 9.06 | 235.1692 | 3,5-Di-tert-butyl-2-hydroxybenzaldehyde | 991 | 1.00 | RPLC | POS |
| 9.13 | 219.1743 | 2,6-Di-tert-butyl-4-(4-morpholinylmethyl)phenol | 786 | 0.72 | RPLC | POS |
| 9.15 | 303.1938 | [7-(2-Hydroxypropan-2-yl)-4a-methyl-1-methylidene-2,3,4,5,6,7,8,8a-octahydronaphthalen-2-yl] acetate | 993 | 0.81 | RPLC | POS |
| 9.25 | 339.1934 | * 15-Deoxy-delta-12,14-prostaglandin J2 | 983 | 0.80 | RPLC | POS |
| 9.36 | 205.0859 | 4-(Butoxycarbonyl)benzoic acid | 989 | 0.88 | RPLC | POS |
| 9.36 | 261.2212 | Pinolenic acid | 781 | 0.76 | RPLC | POS |
| 9.42 | 149.0232 | N-(2-Furylmethyl)-1,3-benzodioxole-5-carboxamide | 995 | 1.00 | RPLC | POS |
| 9.42 | 315.1944 | * Aldosterone | 757 | 0.91 | RPLC | POS |
| 9.42 | 337.1777 | * all-trans-4-Ketoretinoic acid | 989 | 0.80 | RPLC | POS |
| 9.42 | 163.0389 | 1(3H)-Isobenzofuranone, 3-hydroxy-7-methoxy- | 906 | 0.80 | RPLC | POS |
| 9.46 | 272.222 | N-Lauroylsarcosine | 1000 | 1.00 | RPLC | POS |
| 9.52 | 301.2148 | 5-Oxo-6E,8Z,11Z,14Z-eicosatetraenoic acid | 981 | 0.81 | RPLC | POS |
| 9.72 | 289.2139 | Epitestosterone | 1000 | 1.00 | RPLC | POS |
| 9.82 | 299.2581 | 8-(3-Octyl-2-oxiranyl)octanoic acid | 785 | 0.80 | RPLC | POS |
| 9.94 | 301.2148 | 20-Hydroxyarachidonic acid | 981 | 0.79 | RPLC | POS |
| 9.96 | 415.2453 | 9-Oxo-11-alpha,16R-dihydroxy-17-cyclobutyl-5Z,13E-dien-1-oic acid | 893 | 0.67 | RPLC | POS |
| 10.11 | 333.2768 | cis-4,10,13,16-Docosatetraenoic acid | 983 | 0.80 | RPLC | POS |
| 10.15 | 365.3039 | * 2-Arachidonyl glycerol ether | 729 | 0.95 | RPLC | POS |
| 10.20 | 235.2056 | Palmitic acid alkyne | 735 | 0.82 | RPLC | POS |
| 10.34 | 300.2898 | D-erythro-N-stearoylsphingosine | 714 | 0.80 | RPLC | POS |
| 10.41 | 431.2788 | Andrastin F | 941 | 0.69 | RPLC | POS |
| 11.00 | 111.044 | 4-Ketopimelic acid | 929 | 0.75 | RPLC | POS |
| 11.01 | 393.2979 | 15(S)-15-Methylprostaglandin F2-alpha isopropyl ester | 729 | 0.81 | RPLC | POS |
| 11.39 | 340.3576 | Docosanamide | 986 | 0.72 | RPLC | POS |
| 11.57 | 688.4913 | 1,2-Dipalmitoleoyl-sn-glycero-3-phosphoethanolamine | 820 | 0.72 | RPLC | POS |
| 11.89 | 716.5231 | 2-Linoleoyl-1-palmitoyl-sn-glycero-3-phosphoethanolamine | 1000 | 1.00 | RPLC | POS |
| 12.17 | 692.5224 | 1,2-Dipalmitoyl-sn-glycero-3-phosphoethanolamine | 1000 | 1.00 | RPLC | POS |
| 12.27 | 718.5384 | 1-Palmitoyl-2-oleoyl-sn-glycero-3-phosphoethanolamine | 982 | 0.79 | RPLC | POS |
| 16.02 | 338.342 | Erucamide | 712 | 0.88 | RPLC | POS |
| 0.57 | 265.1475 | Laurylsulfuric acid | 961 | 0.70 | RPLC | NEG |
| 0.93 | 173.0093 | cis-Aconitic acid | 994 | 1.00 | RPLC | NEG |
| 1.50 | 195.051 | D-Gluconic acid | 991 | 0.95 | RPLC | NEG |
| 1.76 | 227.056 | 4-Nitro-N-(2-pyrimidinyl)benzamide | 940 | 0.76 | RPLC | NEG |
| 1.84 | 306.076 | S-Lactoylglutathione | 938 | 0.84 | RPLC | NEG |
| 2.05 | 178.0511 | 1-(3-Pyridyl)-1-butanone-4-carboxylic acid | 999 | 1.00 | RPLC | NEG |
| 2.22 | 101.0245 | (2E)-4-Hydroxybut-2-enoic acid | 959 | 0.96 | RPLC | NEG |
| 2.36 | 130.0511 | Acetyl-beta-alanine | 957 | 1.00 | RPLC | NEG |
| 2.70 | 131.0715 | 2-Ethyl-2-hydroxybutyric acid | 1000 | 1.00 | RPLC | NEG |
| 3.01 | 161.0458 | beta-D-Glucopyranoside, 4-[(4-carboxy-3-hydroxy-3-methyl-1-oxobutoxy)methyl]phenyl | 965 | 0.73 | RPLC | NEG |
| 3.09 | 176.0388 | N-Formyl-L-methionine | 863 | 0.68 | RPLC | NEG |
| 3.58 | 212.0023 | Indoxyl sulfate | 989 | 0.80 | RPLC | NEG |
| 4.01 | 188.0566 | N-Acetyl-L-glutamic acid | 996 | 1.00 | RPLC | NEG |
| 5.25 | 103.0402 | 3-Hydroxybutyric acid | 1000 | 1.00 | RPLC | NEG |
| 6.07 | 252.0697 | N-Acetyl-S-benzyl-L-cysteine | 983 | 0.80 | RPLC | NEG |
| 8.95 | 159.0301 | 2-Oxohexanedioic acid | 887 | 0.67 | RPLC | NEG |
| 1.26 | 255.0975 | 5-Methoxy-1H-benzo[d]imidazole-2-carboxylic acid | 856 | 0.89 | HILIC | POS |
| 1.26 | 195.1128 | (5aR,10aR)-Octahydrodipyrrolo[1,2-a:1',2'-d]pyrazine-5,10-dione | 1000 | 1.00 | HILIC | POS |
| 1.38 | 100.0756 | 1-Piperidinecarboxylic acid, 2-oxo-, 1,1-dimethylethyl ester | 829 | 0.76 | HILIC | POS |
| 1.49 | 123.0552 | N-(Hydroxymethyl)nicotinamide | 937 | 0.99 | HILIC | POS |
| 1.75 | 193.1699 | (S)-Amiflamine | 821 | 0.80 | HILIC | POS |
| 2.06 | 135.0804 | 3-Isobutylpentanedioic acid | 973 | 0.97 | HILIC | POS |
| 2.35 | 290.1387 | * Norcocaine | 996 | 1.00 | HILIC | POS |
| 4.05 | 104.1069 | Glycerophosphocholine | 946 | 0.84 | HILIC | POS |
| 4.98 | 112.0505 | Cytidine 3'-monophosphate | 924 | 0.83 | HILIC | POS |
| 5.49 | 137.0709 | 1-Methylnicotinamide cation | 883 | 0.88 | HILIC | POS |
| 7.04 | 118.0862 | Betaine | 1000 | 1.00 | HILIC | POS |
| 0.96 | 213.0557 | 2,4'-Dihydroxybenzophenone | 910 | 0.84 | HILIC | NEG |
| 1.00 | 121.0297 | 5-Formyl-2-hydroxybenzene-1-sulfonic acid | 995 | 0.82 | HILIC | NEG |
| 1.08 | 229.0868 | 3-Hydroxy-5-[2-(4-hydroxyphenyl)ethyl]phenyl-beta-D-glucopyranoside | 1000 | 1.00 | HILIC | NEG |
| 1.11 | 175.025 | * Vitamin C | 870 | 0.76 | HILIC | NEG |
| 1.16 | 151.0402 | 3-Methoxybenzoic acid | 1000 | 1.00 | HILIC | NEG |
| 1.17 | 200.0023 | 3-Sulfamoylbenzoic acid | 987 | 0.72 | HILIC | NEG |
| 1.28 | 259.0972 | 5-Homovanillylresorcinol | 974 | 0.79 | HILIC | NEG |
| 2.60 | 261.0071 | * Homovanillic acid sulfate | 976 | 0.91 | HILIC | NEG |
| 3.49 | 222.0771 | N-Acetyl-L-tyrosine | 961 | 0.83 | HILIC | NEG |
| 4.52 | 157.0369 | Allantoic acid | 761 | 0.98 | HILIC | NEG |

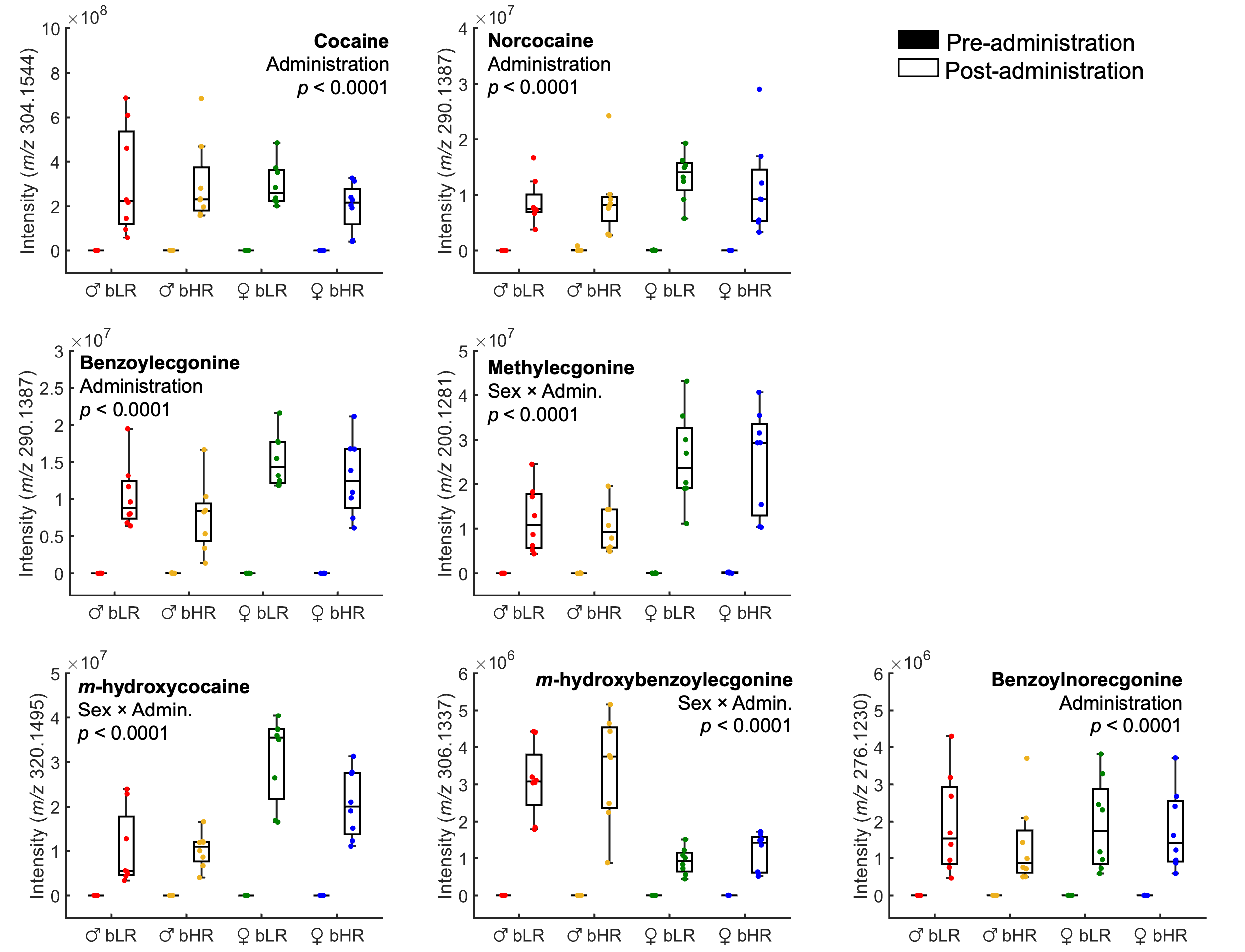

**Figure S6.** Cocaine metabolites measured in brain dialysate. Box plots are shown with center line as median, extending to the IQR, with whiskers to the highest and lowest non-outlier points. Each dot represents data from a single animal (*N* = 8 rats/group). Calculated *p*-values for either the main effect of drug administration or sex × administration interaction are also indicated. Post hoc analysis revealed that extracellular levels of methylecgonine and *m-*hydroxycocaine were higher in females while *m*-hydroxybenzoylecgonine levels were higher in males.

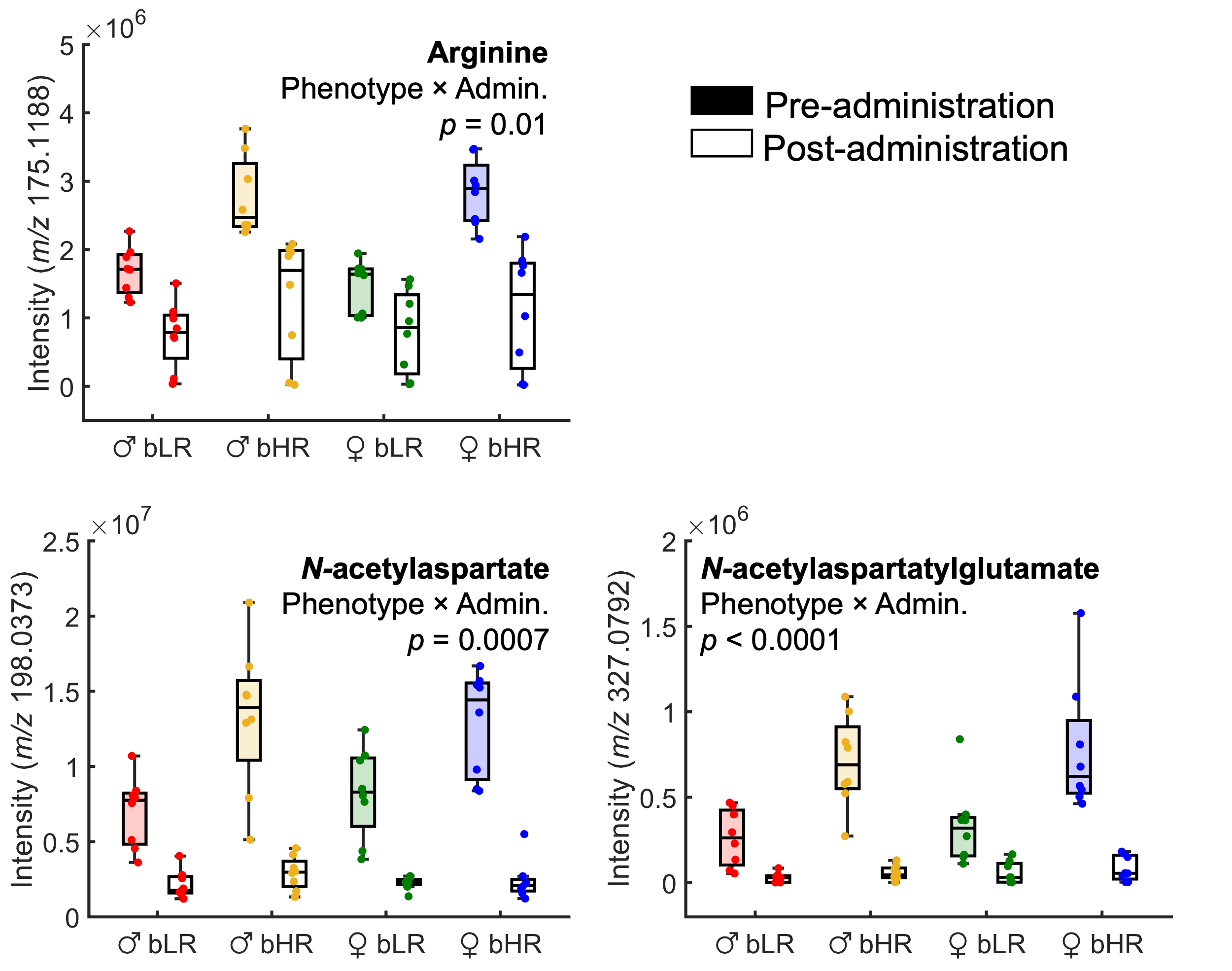

**Figure S7.** Changes in metabolites linked to arginine biosynthesis and alanine, aspartate, and glutamate metabolism following cocaine administration. Box plots are shown with center line as median, extending to the IQR, with whiskers to the highest and lowest non-outlier points. Each dot represents data from a single animal (*N* = 8 rats/group). Calculated *p*-values for phenotype × administration interaction are provided. Post hoc testing showed no significant phenotypic differences for these compounds after administration.
